# Two common and distinct forms of variation in human functional brain networks

**DOI:** 10.1101/2021.09.17.460799

**Authors:** Ally Dworetsky, Benjamin A. Seitzman, Babatunde Adeyemo, Ashley N. Nielsen, Alexander S. Hatoum, Derek M. Smith, Thomas E. Nichols, Maital Neta, Steven E. Petersen, Caterina Gratton

## Abstract

The cortex has a characteristic layout with specialized functional areas forming distributed large-scale networks. However, substantial work shows striking variation in this organization across people, which relates to differences in behavior. While most prior work treats all individual differences as equivalent and primarily linked to boundary shifts between the borders of regions, here we show that cortical ‘variants’ actually occur in two different forms. In addition to border shifts, variants also occur at a distance from their typical position, forming ectopic intrusions. Both forms of variants are common across individuals, but the forms differ in their location, network associations, and activations during tasks, patterns that replicate across datasets and methods of definition. Border shift variants also track significantly more with shared genetics than ectopic variants, suggesting a closer link between ectopic variants and environmental influences. Further, variant properties are categorically different between subgroups of individuals. Exploratory evidence suggests that variants can predict individual differences in behavior, but the two forms differ in which behavioral phenotypes they predict. This work argues that individual differences in brain organization commonly occur in two dissociable forms – border shifts and ectopic intrusions – suggesting that these types of variation are indexing distinct forms of cortical variation that must be separately accounted for in the analysis of cortical systems across people. This work expands our knowledge of cortical variation in humans and helps reconceptualize the discussion of how cortical systems variability arises and links to individual differences in cognition and behavior.

## 1. INTRODUCTION

The cortex shows a characteristic organization, with distinct functional areas linking together to form distributed large-scale systems (i.e., networks^1^). This organization follows a stereotyped general pattern, but is not completely uniform, varying both across and within species. Comparative neuroanatomy studies demonstrate that variations in cortical organization appear in a constrained set of possible forms, including changes in the relative size, connectivity, or functional characteristics of cortical fields^2, 3^. Notably, similar variation in cortical organization also exists within a species and can be influenced by genetic and developmental factors^4, 5^. Variation in cortical organization relates to differences in phenotypic characteristics and behavior^2^, suggesting that studying variation in cortical functional architecture in humans may provide insights into the sources of varying behavioral traits relevant to cognition and disease.

The regional and systems-level organization of the human brain can be mapped non-invasively using functional connectivity MRI (*fcMRI;* correlations in the spontaneous activity patterns between different regions). FcMRI can be used to identify functionally homogenous regions^6–8^ and distinct systems^9, 10^ that correspond well to patterns detected with task activation methods. For large groups of individuals, a “typical” or “canonical” average pattern of distributed functional systems emerges that is reproducible across studies and maps onto differences in motor, sensory, and higher-level processing^9, 10^.

However, recent work has also highlighted that any given person differs from this group pattern, at least in some locations^11–20^. We characterized locations, that we call *network variants*, where an individual’s functional connectivity pattern differs markedly from the typical group average (with similarity below r<0.3)^15, FN1^. Across multiple datasets, we demonstrate that network variants are stable over time and across task states^21^ and relate to individual differences in task-related brain activations and behavioral measures collected outside of the scanner^15^. Moreover, different subgroups of individuals show similarities in their network variants: one subgroup of individuals shows network variants more associated with top-down control and sensorimotor systems, and another subgroup shows network variants more associated with the default mode system. We hypothesize that network variants reflect trait-like variations in the organization of functional brain areas across individuals^15^.

While a number of studies have identified locations of individual differences in brain network organization^11–20^, there has not yet been substantial research into the different forms that these variants take and how they might link to sources and mechanisms observed in past comparative studies of cortical neuroanatomy in other species^2^. Many studies, including our past work, treat all forms of cortical variation uniformly, and often assume that variants are driven primarily by boundary shifts in the borders between systems: a functional region may expand, contract, or be slightly offset relative to the typical pattern (e.g., as has been documented for V1, which can differ more than 2-fold in size across individuals^22, 23^). These *border shifts* are likely important for understanding individual differences in behavior and brain function^19, 24–27^, and could potentially be addressed through functional alignment methods that allow for individual-specific regions to be locally displaced (e.g., refs.^24, 28^).

However, it is possible that brain organization can also differ more markedly from the stereotypical pattern, with islands of idiosyncratic fcMRI occurring in locations remote from their typical system organization^29^. We call these shifts in brain organization *ectopic intrusions* to reflect the presence of these variations at abnormal locations (see Fig. 1). Evidence of dramatic shifts in the function or connectivity of primary sensory/motor areas can be seen with systematic deprivations in development^5, 30, 31^, but it is less well understood how common these shifts are in higher-level association areas in neurotypical humans – despite some prior observations that they occur^11, 15^. Ectopic intrusions are not easily explained by cortical expansion mechanisms and will be poorly addressed by functional alignment techniques that assume only local displacements in brain architecture, confounding group studies.

**Fig. 1:**
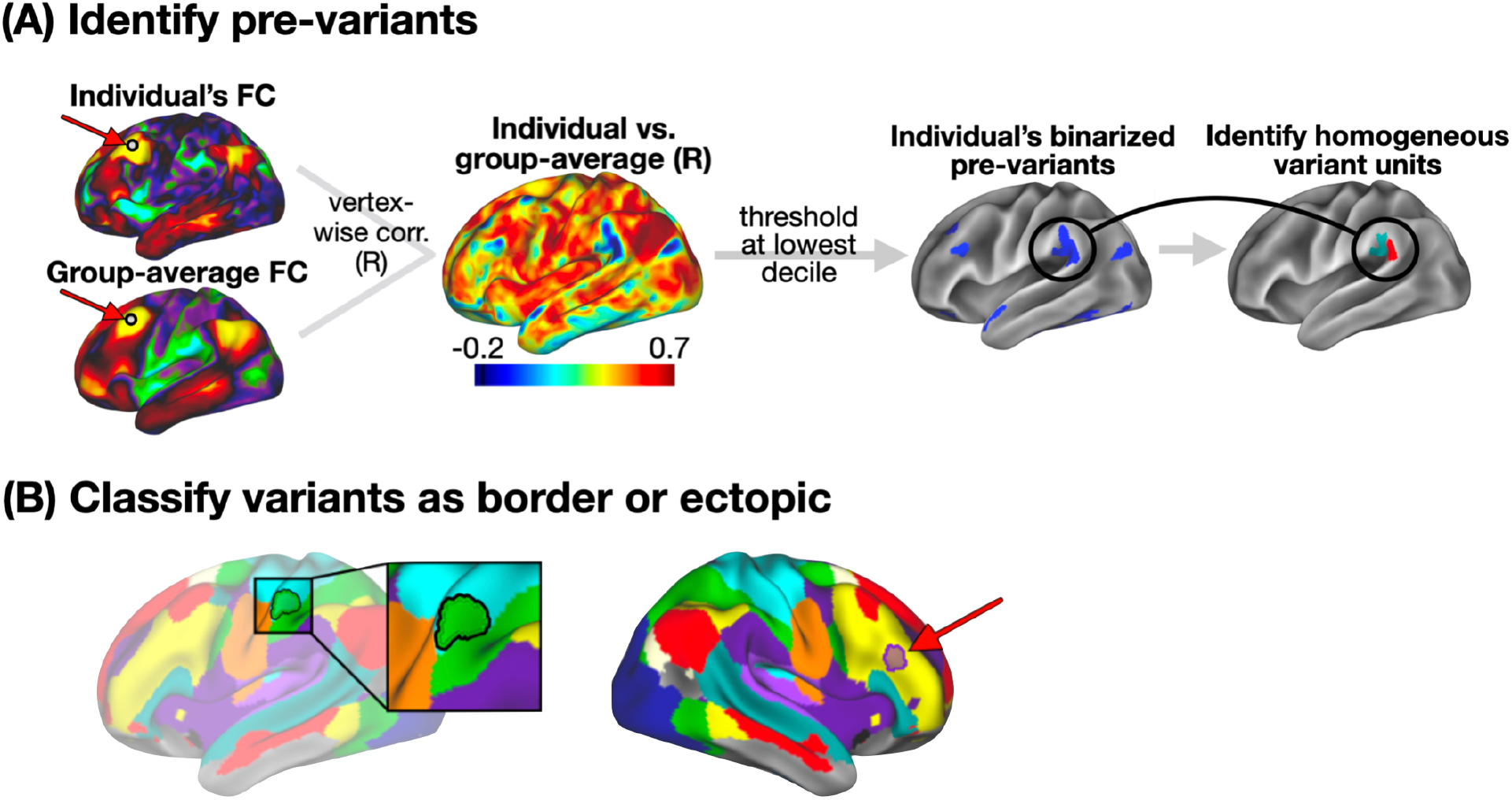
Variant definition, splitting, and classification as border or ectopic. (A) Following Seitzman et al.^15^, we used spatial correlation to compare the seedmap at a given location between an individual and an independent group average (left) to generate an individual-to-group “similarity map” (middle). This similarity map was thresholded and binarized to identify locations with low similarity to the group (right) that we call “pre-variants” in this work (Note: these were also thresholded to remove small areas and areas of low signal - see *Methods*). We then further refined these pre-variants to create homogeneous units for border vs. ectopic variant classification (see Methods and Supp. Fig. 1 for a description of this process). (B) Each variant was classified as either a border shift or an ectopic intrusion based on its edge-to-edge distance from the nearest same-network boundary in the group-average network map. Here, we display an example of a border shift variant (left, green region with black outline) and an ectopic variant (right, purple region with dark outline), overlayed on the group average network map. Distances > 3.5 mm were classified as ectopic; distances < 3.5 mm were classified as border (see Fig. 2B and Supp. Fig. 8 for exploration of additional distance criteria). We also employed a secondary method for defining border and ectopic variants that did not rely on a group-level network parcellation (see description in *Methods* and Supp. Fig. 2).

The goal of this study was to examine the relative prevalence of ectopic intrusions and border shift variants in humans, and to contrast the properties of these two forms of idiosyncratic variation. To this end, we used a combination of data from a highly sampled “precision” fMRI dataset and a larger-*N* dataset to assess how commonly each of these forms of individual differences are present across people and the extent to which they vary in their characteristics. Separating these forms will likely be essential to deepening our understanding of the sources and consequences of individual differences in human brain organization and their relevance to clinical disorders.

## 2. RESULTS

Using a combination of the large HCP dataset (N = 374 unrelated individuals used with at least 40 min. of data used for primary analyses, and N = 793 individuals used for analyses of behavior and similarity among twin samples) and the precision MSC dataset (N = 9, scanned 10 times each), we investigated two different forms of variation in functional system organization: variations associated with nearby shifts in the boundaries between a person’s network organization and the typical layout (“border” shift variants) and more remote islands of functional networks not adjacent to their typical layout (“ectopic” intrusion variants; see Fig. 1 and *Methods* for information on how both forms of variants are defined). The two datasets were used in combination to leverage their relative strengths in terms of sample size and data quantity per individual, respectively, and to demonstrate replicability of this work.

In our analyses, we first examined how common ectopic intrusions were relative to border shifts variants across participants. We next asked how the two variant forms compare in their spatial occurrence, the networks they are affiliated with, and how they respond during tasks. We contrasted variants in familial samples to estimate how they are affected by genetic similarity, and we examined whether subgroups of individuals showed similar forms of border and ectopic variants. Finally, we examine how border and ectopic variants predict behavioral phenotypes collected outside of the scanner.

### 2.1. Nearly all individuals exhibit both border and ectopic cortical variants

Our first goal was to establish how frequently each of these forms of network variants occur. We used two different methods for defining border shifts and ectopic intrusions. Our primary method (Fig. 1, *Methods*) defines border shifts and ectopic intrusions based on their distance to pre-defined canonical (i.e., previously published group-average) network boundaries^11^. Our secondary method (reported in the supplement; see Supp. Fig. 2 for schematic) used a parcellation-free approach to define these two forms of variants, based on a continuous measure of similarity to nearby locations in the group-average (i.e., finding a location with 90% maximum similarity within 10mm.). This approach has the advantage that it does not depend on a specific group parcellation for networks (e.g., group networks from ref. ^11^ vs. ref. ^10^) or network resolution (e.g., 7 vs. 17 networks from Yeo et al.^10^).

Using either method, both ectopic and border variants were consistently identified in almost all individuals in both the MSC and the HCP datasets: at least one ectopic variant and at least one border variant were observed in all 9 of 9 MSC subjects and in 371 of 374 HCP subjects (> 99%). In the MSC, an average of 49.9% [±10.4%] of an individual’s network variants were ectopic; in the HCP, an average of 42.6% [±14.9%] of an individual’s network variants were ectopic (Fig. 2A; see Supp. Fig. 3 for similar results with secondary method). As can be seen in Fig. 2A (see also Supp. Fig. 4), the specific proportion of border and ectopic variants differed somewhat across individuals, but the proportion of ectopic variants did not correlate with the total number of variants within an individual (r = 0.15 in the MSC dataset and r = 0.05 in the HCP). Interestingly, increasing the distance criteria (from 3.5 to 5, 7.5, or 10 mm) still left a large proportion of ectopic variants, with roughly 30% of variants classified as ectopic at 10 mm (Fig. 2B). Indeed, the median distance between ectopic variants and their own network was > 15 mm (Supp. Fig. 5). These results are consistent with findings from our secondary method for defining border and ectopic variants (see Supp Fig. 6). Note that similar results are also obtained when ectopic variants are defined relative to a participant’s own network boundaries (see Section 5.5 for description of methods, Supp. Fig. 7 and Supp. Table. 1). Thus, distant ectopic variants are not rare phenomena and, along with border variants, appear to be relatively ubiquitous at the individual level.

**Fig. 2:**
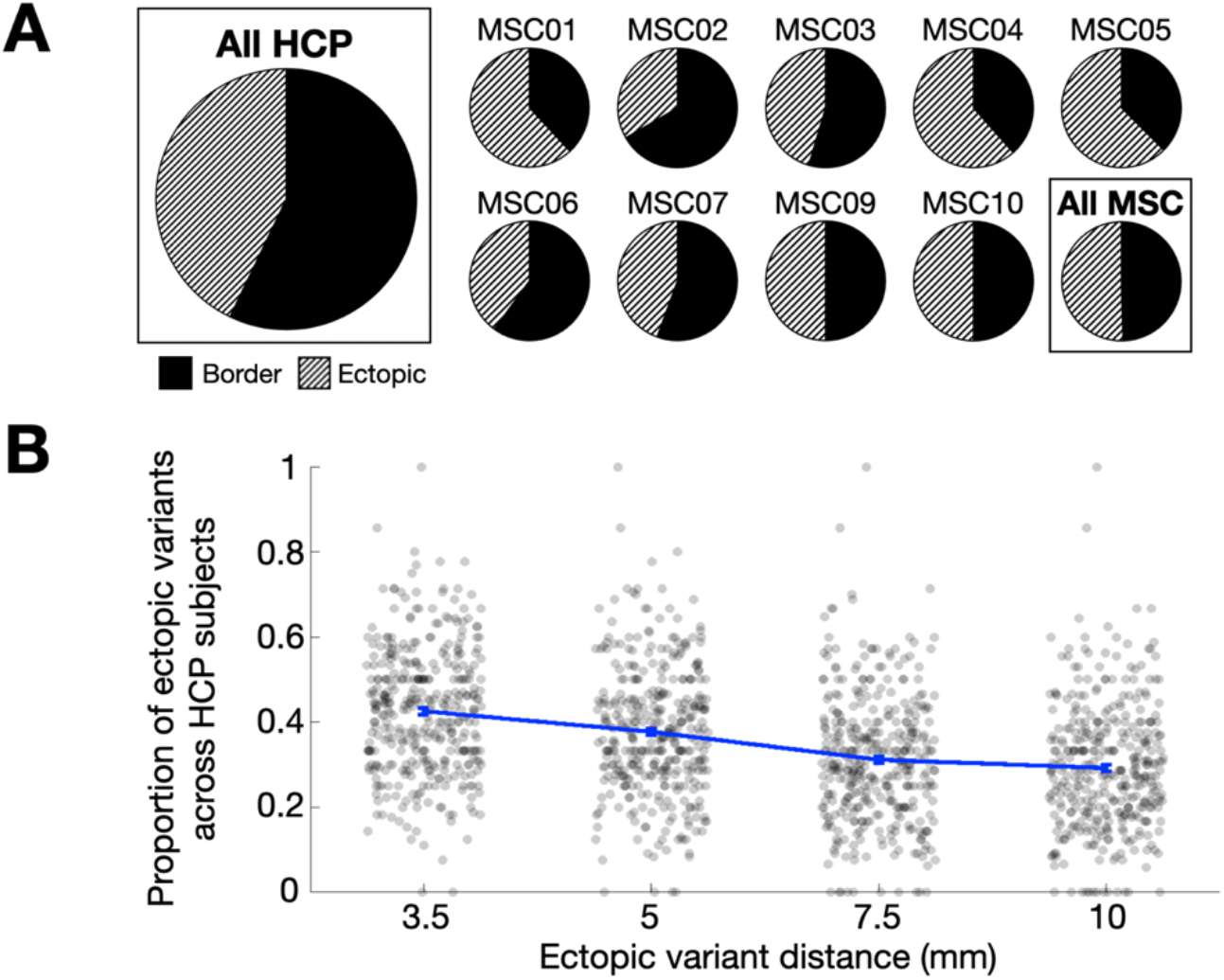
Prevalence of border and ectopic variants across individuals. (A) The panel displays the proportion of border and ectopic variants across all subjects in the HCP dataset (far left), and within subjects in the MSC dataset (two rows on right). Both ectopic variants and border variants were consistently identified in almost all individuals in both the HCP and the MSC datasets. (B) Proportions of ectopic variants at other distances (error bars represent SEM across participants; each dot represents a single subject). As the required minimum distance for a variant to be classified as ectopic increases to 5, 7.5, and 10 mm, ectopic variants continue to comprise a sizable percentage of all variants in the HCP dataset, nearly 30%, even at a distance of 10 mm. Indeed, the median distance for ectopic variants from a network border was > 15 mm (Supp. Fig. 5). Similar results are seen with our secondary parcellation-free method of defining network variants (Supp. Fig. 3, 6).

### 2.2. Border and ectopic variants show significant differences in their spatial distribution

Next, we asked whether ectopic and border variants tend to occur in different locations. We examined the spatial overlap of ectopic variants and compared it to the overlap of border variants. The spatial distributions of the two variant forms are shown in Fig. 3A, where warmer colors represent brain regions with a high occurrence of variants across subjects. As a general pattern, both forms of variants follow the distribution previously described of higher prevalence in association regions of cortex^16, 17, 19^, especially in lateral frontal cortex, superior frontal cortex and near the temporoparietal junction^15^. However, direct contrasts between the two forms of variants suggest that ectopic variants appear more frequently in some locations than border shifts, and vice versa. Notably, border and ectopic variant spatial distributions were similar between the HCP and MSC datasets; see Supp. Fig. 8.

**Fig. 3:**
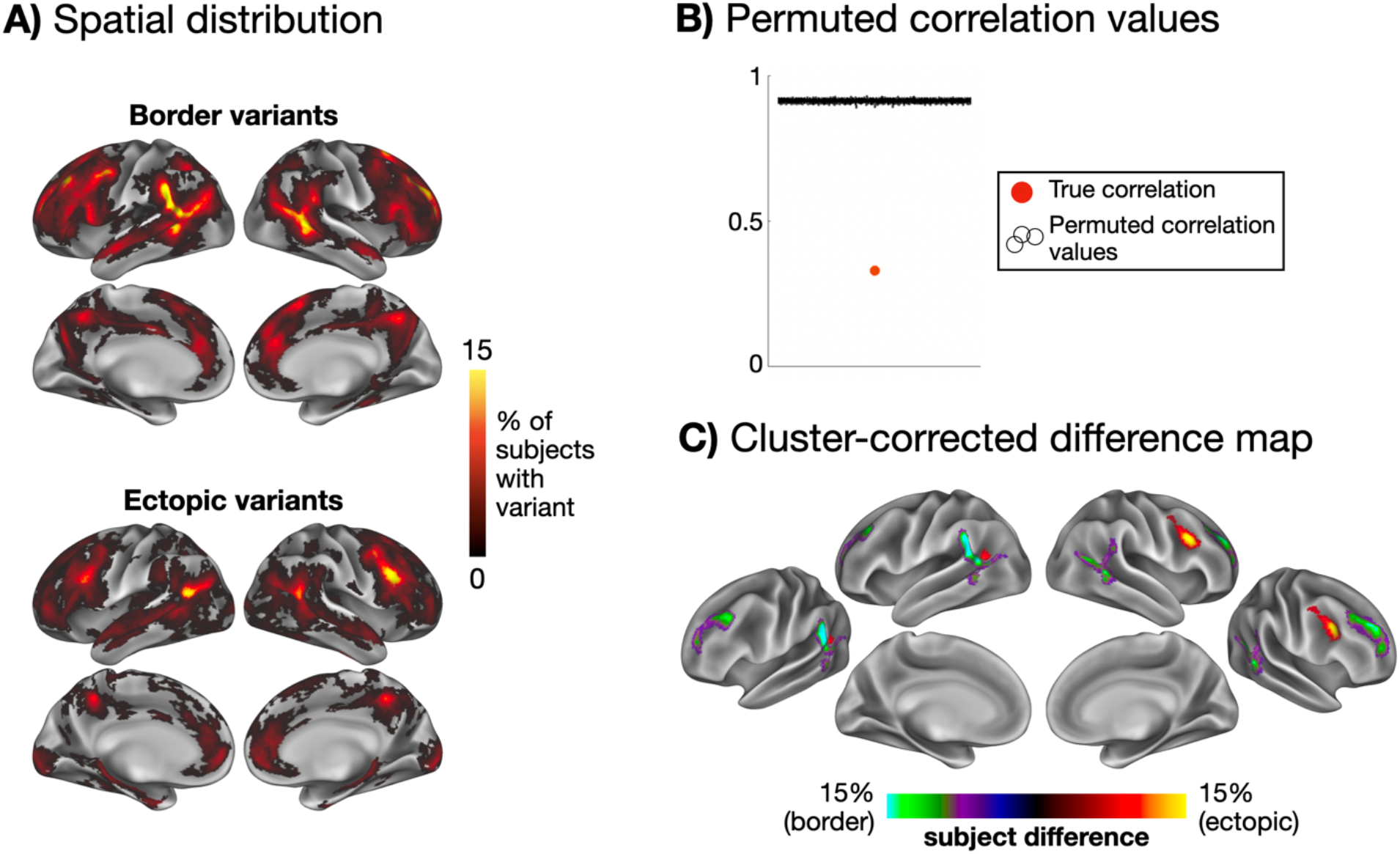
Spatial distributions of ectopic and border variants. The maps in (A) show the spatial distribution of border variants and ectopic variants, overlapped across participants in the HCP. (B) These two spatial distributions differ significantly more than expected by chance from random permutations. Permuted correlation values are jittered across the x-axis for visualization. (C) A cluster-corrected difference map is shown highlighting regions with a significantly higher occurrence of border variants (green/purple) and ectopic variants (yellow/red; p < 0.05 cluster-corrected for multiple comparisons based on permutation testing). Ectopic variants were more prevalent in the right posterior inferior frontal sulcus and left posterior TPJ regions, while border variants were more prevalent in dorsal and ventral portions of the anterior TPJ and superior rostral frontal regions. See Supp. Fig. 9 for evidence of similar results for ectopic variants at greater distances, Supp. Fig. 10 for similar results using our secondary parcellation-free variant definition method, and Supp. Fig. 11 for border and ectopic overlap maps for each network individually.

To assess the significance of these differences, we conducted a permutation test where ectopic and border labels were randomly flipped within subjects and used to create 1000 permuted overlap maps. The true maps were then compared to these permuted maps. The true ectopic and border variant maps were significantly more dissimilar than the permuted maps (p < 0.001; none of the permuted maps were as dissimilar as the true overlap maps, Fig. 3B). A similar permutation approach was used to create a multiple comparisons cluster-corrected difference map (p < 0.05, see *Methods*), revealing several key locations that differed in the frequency with which the two variant forms appeared. As shown in Fig. 3C, ectopic variants appear more frequently in dorsolateral frontal regions of the right hemisphere, whereas border variants are more frequent around rostral portions of temporoparietal junction and superior rostral frontal regions in both hemispheres (see Supp. Fig. 9 for similar results with ectopic variants defined at longer distances, Supp Fig. 10 for similar results with our secondary parcellation-free definition method).

### 2.3. Border and ectopic variants exhibit different patterns of network assignment

We next examined the network associations of each form of variant. As described in the *Methods*, variants were “assigned” to a network by identifying the template (see Supp. Fig. 12) that best fit that variant’s seedmap^FN2^. As a general pattern, variants tend to frequently be assigned to the default mode, fronto-parietal, and cingulo-opercular networks for both forms and across both datasets, consistent with previously reported results (Fig. 4A, Supp. Fig. 13). However, when contrasted with one another, we find that the relative prevalence of border and ectopic variants differs by network. Indeed, a mixed-effects generalized linear model analysis revealed a significant interaction between variant form and network (p<0.001), suggesting that the influence of variant form on subject-level variant frequencies varies significantly across networks.

**Fig. 4:**
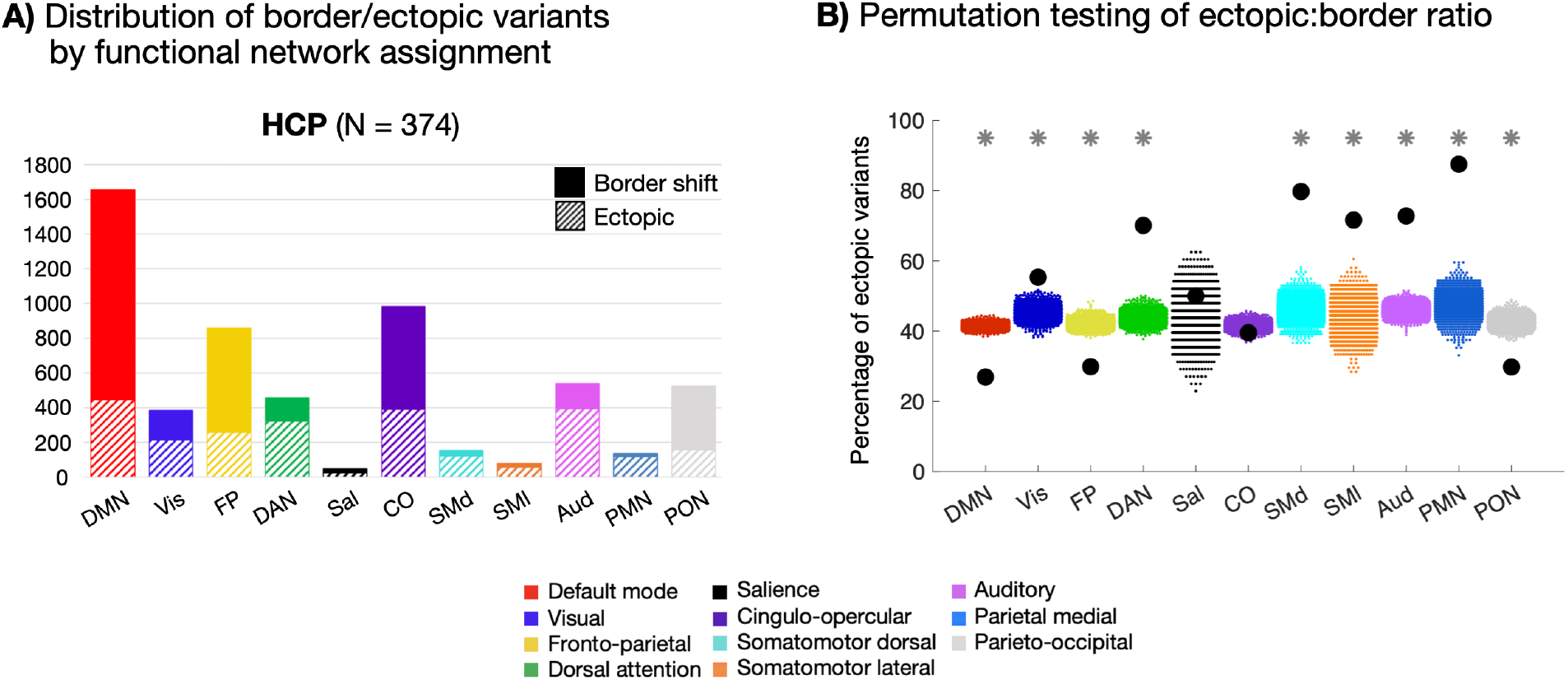
Network linkages of border and ectopic variants. (A) Network distributions of border and ectopic variants in the HCP dataset. Variants of both forms are commonly associated with the DMN, FP, and CO networks, as reported in past work^15^. Similar results were seen in the MSC dataset (Supp. Fig. 13) and using the parcellation-free approach to defining border and ectopic variants (Supp. Fig. 14) (B) Plot depicting permutation testing of the ectopic:border ratio in the HCP dataset. For all networks with the exception of salience and cingulo-opercular, the true proportion of ectopic variants (black dots) was significantly different from permuted proportions (colored dots, 1000 random permutations of shuffled labels) at *p*<0.001 (*; FDR corrected for multiple comparisons). DMN, FP, and PON variants were more likely to be border shifts, while sensorimotor, DAN, and PMN variants were more likely to be ectopic. Notably, ectopic variants were commonly found in all systems. See Supp. Fig. 12 for cortical depiction of each listed network.

Using our primary variant definition method in the HCP dataset, border variants appeared more commonly assigned to the default mode network (DMN), fronto-parietal (FP), and parieto-occipital network (PON). Ectopic variants were relatively more linked to the dorsal attention (DAN), parietal memory (PMN), and sensorimotor networks (visual, somatomotor, auditory; see also similar results using our secondary method, Supp. Fig. 14). Permutation testing of variant labels in the HCP dataset confirmed these observations (Fig. 4B). As may be expected, ectopic variants were relatively more abundant in smaller or more spatially local networks, as it is easier to be distant from these network boundaries. However, ectopic variants were still often associated with large networks such as the DMN, FP, cingulo-opercular (CO), DAN, and Visual systems, emphasizing their common nature. Distributions of variant network assignments in the MSC are shown in Supp. Fig. 13. Although there were fewer variants in the MSC overall (leading to greater variability in per-network proportions of variant forms), the findings are consistent with the HCP dataset such that ectopic variants were associated with many large (and small) networks. The parcellation-free classification method also resulted in comparable network distributions for most networks, although with some differences in ectopic:border ratios (Supp. Fig. 14).

Interestingly, combining information from the location and network-assignment analysis, one can find that border and ectopic variants exhibited different patterns of “swaps” in their territory relative to the canonical structure (see Supp. Fig. 15). For example, border variants found in canonical (i.e., group average) regions of the FP network most often are re-assigned to DMN, CO, or DAN. Thus, while idiosyncratic, both forms of variants show constraints in how they vary.

While it is beyond the scope of this manuscript to fully explore the interactions between variant form (border, ectopic) and network assignment, it will be useful in future work to examine this question in more detail. To aid in this endeavor, in ***Supp. Fig. 11*** we provide border and ectopic variant maps for each network individually.

### 2.4. Border and ectopic variants exhibit shifted task responses

Next, using both datasets, we asked if both forms of variants show altered task responses, consistent with their idiosyncratic network affiliation. Following Seitzman et al.^15^, we first focused on the mixed design task from the MSC. This design consisted of a cued-block paradigm, with blocks of noun/verb judgments on presented words and blocks of dot coherence concentricity judgments. Average activations (for all cue/trial/block conditions) were contrasted with baseline (see *Methods* for details), a contrast that elicits negative activations in DMN regions typically, and positive activations in the FP, DAN, and visual systems^15, 32^.

In the MSC, ectopic variants and border variants exhibit shifted task activation responses, in the direction expected for canonical regions associated of the same network (Fig. 5B). Border variants are shifted further toward the expected activation based on their new assignment than ectopic variants. While we observe a strong task deactivation for border DMN variants (t(7) = −5.38, *p* = 0.001 for 8 MSC participants with border DMN variants), aligning closely with canonical DMN deactivation, ectopic DMN variants exhibit a relatively weak (but still significant) deactivation (t(6) = −2.76, *p* = 0.033 for 7 MSC participants with ectopic DMN variants). Similar patterns are seen with “task positive” networks like the visual, FP, and DAN networks; the CO, language, and salience networks were not strongly modulated by this contrast. This finding is further examined in Fig. 5A for each MSC participant by comparing the task activation of DMN variants in a given subject to that same location in other subjects. This analysis showed that – in every individual participant – similar patterns of prominent decreases in activations were observed relative to what is expected for that location in both border variants (t(7) = 5.97, p<0.001) and ectopic variants (t(6) = 5.99, p<0.001). However, the ectopic DMN variants appear in locations that have a more positive typical response pattern in comparison to border variants, and thus do not reach as strong a level of deactivation. We again found similar results with the secondary parcellation-free classification method (see Supp. Fig. 16).

**Fig. 5:**
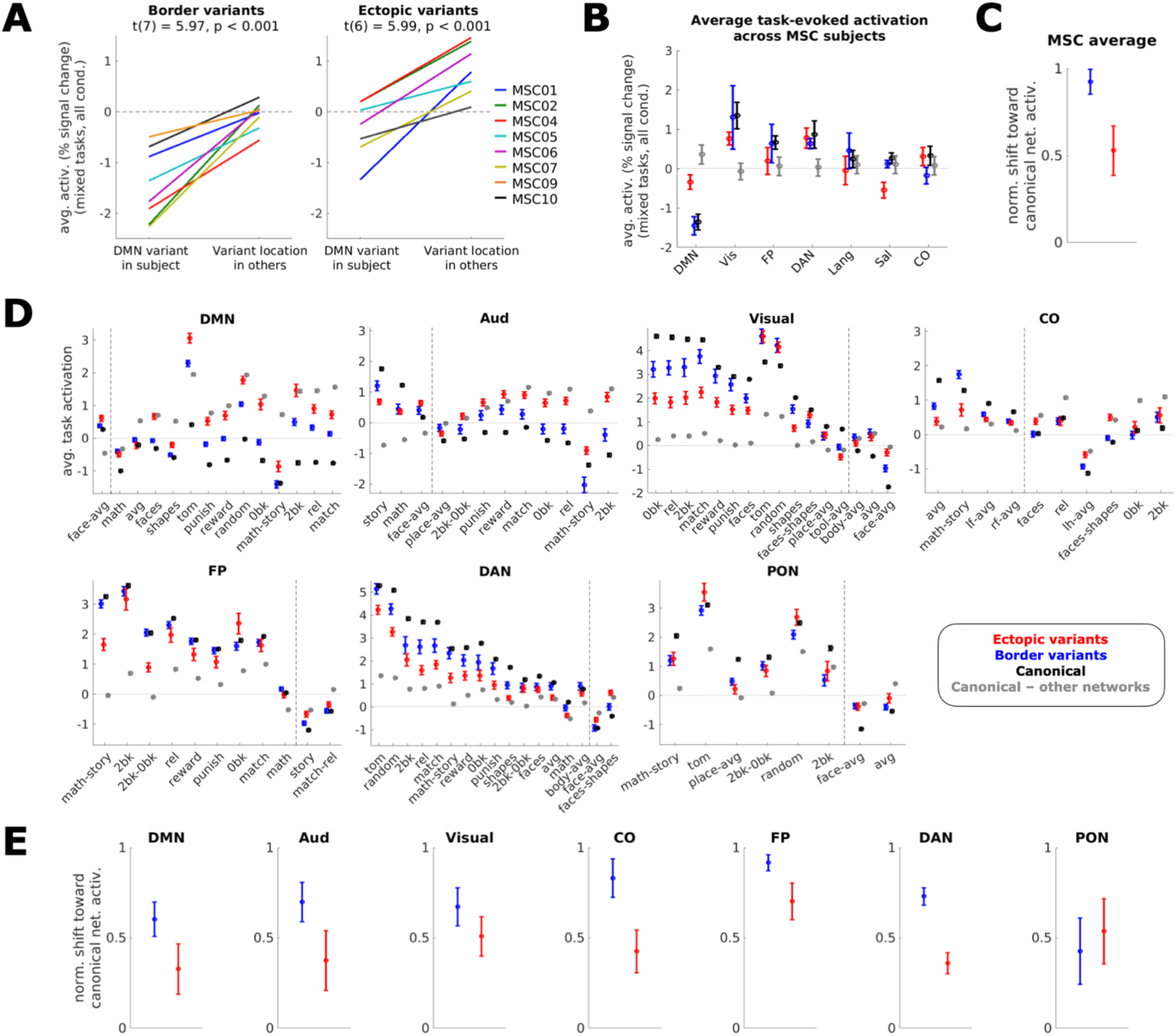
Task-evoked activation of variant locations in the MSC and HCP. (A) Average activation in the MSC of DMN-assigned variants in an individual vs. average activation in that location in all other individuals for border (left) and ectopic (right) variants; 8 of 9 subjects with a border DMN variant and 7 of 9 subjects with an ectopic DMN variant were included. Different colors represent different MSC participants. Both border and ectopic variants exhibited functional activation shifted from the typical responses of their location toward the response expected based on their network associations. Similar results were seen with our secondary method of border/ectopic variant classification (Supp. Fig. 16). (B) Average activation (*z*) across all task conditions in a set of mixed-design tasks from the MSC for variants (red = ectopic, blue = border shift), canonical locations of the listed network (black), or canonical locations of other networks (gray). Error bars represent the standard error of the mean across subjects. Data were only included from individuals that had a variant of the given form and network (e.g., a border variant assigned to the DMN). (C) Normalized shift of border (blue) and ectopic (red) variants toward the listed canonical networks. Values represent the average normalized value across the listed networks (calculated as *(variant–other networks)÷(canonical network–other networks)*); error bars represent standard error of the mean across networks. (D) Average task activation across HCP subjects. For each network, contrasts were included if the activation in canonical regions of the network was greater than that of other networks by a margin of at least 0.5% signal change. Vertical dashed lines mark the delineation between contrasts where a given canonical network’s activation is greater than other networks’ activation and contrasts where other networks’ activation is greater. Error bars represent standard error of the mean across subjects for a given contrast; see Supp. Table 2 for information on tasks associated with each contrast name. (E) Normalized shift of border (blue) and ectopic (red) variants toward each listed network. Similarly to the MSC dataset, both border and ectopic variants exhibit shifted responses during tasks. Border variants consistently shift more strongly toward activation of canonical regions of their assigned network than ectopic variants do (with the exception of PON variants). Error bars for the normalized values represent standard error of the mean across contrast activation values within a network.

Next, we turned to the HCP to extend these results to a dataset with a larger sample size. Analysis-level task fMRI maps from the HCP dataset (MSM-Sulc registered, 4mm-smoothing versions of publicly available data) were used to query all contrasts across the 7 tasks (emotional processing, gambling, language, motor, relational processing, social cognition, and working memory). As in the MSC, we compared task activations at variant locations with activations of canonical regions of a variant’s assigned network, and activations of canonical regions of all other networks. Using 358 of 374 subjects with both forms of variants and task data available for all HCP tasks, Fig. 5D shows these results for each contrast by network (including any contrast for which the given network’s canonical response was at least 0.5% higher/lower than *other* networks’ response). See Supp. Table 2 for information on the contrast names by task. Again, border variants are shifted further toward the expected direction for canonical regions of the same network than are ectopic variants (Fig. 5D). The consistency of this result was confirmed by calculating the activation shift of the border or ectopic variant toward the response of regions of its canonical network, normalized by the response of regions of all *other* networks (Fig. 5E). The same analysis was also applied to the MSC task results for consistency (Fig. 5C). In all cases in both datasets, border and ectopic variants both exhibited shifts in their task activations toward canonical network responses. For all but one network in the HCP, border variants consistently shift more strongly toward their network’s canonical response than ectopic variants do.

Jointly, these findings demonstrate that both ectopic and border variants are associated with shifted task activations, shifting toward the expected activations for the variant’s assigned network. Border variants associated with the default mode network retain a task activation similar to canonical locations of their assigned network, while ectopic variants show an altered but intermediate task activation response.

### 2.5. Border and ectopic variants both show a genetic influence, but border variants are significantly more similar among identical twins

To better understand what factors contribute to the formation of network variants, we examined their similarity in twin samples using an expanded subset of HCP subjects, now including those with familial relationships. We measured the similarity in location of network variants (see Fig. 6A for schematic) among monozygotic (N = 88 pairs) and dizygotic twins (N = 45 pairs), siblings (N = 137 pairs), and randomly matched unrelated individuals (N = 122 pairs). As can be seen, network variants were generally most similar in location among monozygotic twins, with intermediate similarity among dizygotic twins and siblings, and with the highest dissimilarity among unrelated individuals (Fig. 6C). Similar twin sample similarity results were also found when using the parcellation-free method to classify variants (see Supp. Fig. 17A). This pattern is indicative of a genetic influence in network variant locations. Indeed, estimates of similarity among twin samples based on Falconer’s formula^33, 34^ were significantly higher for both border and ectopic variants relative to permuted null distributions (p<0.001 and p<0.002 respectively; Fig. 6B). These permutation test results also replicated using the parcellation-free classification methods (p<0.001 for both border and ectopic variants; see Supp. Fig. 17B). These findings are consistent with past reports that functional network organization is heritable^4, 35^, but here are extended to demonstrate genetic influence specifically for the locations of idiosyncratic areas (variants) of the brain.

Intriguingly, when border and ectopic variants are compared with one another, border variants appear to be more similar across identical twin pairs than ectopic variants (Fig. 6C). A two-way mixed-effects ANOVA confirmed this impression, revealing a significant interaction between variant form and group (Greenhouse-Geisser-corrected p<0.001). Unpaired t-tests (assuming unequal variance) were performed to decompose this interaction, demonstrating that monozygotic twins show a greater difference between border and ectopic variant similarity compared to dizygotic and non-twin sibling pairs (p=0.015; significant after Bonferroni correction), and to unrelated individuals (p<0.001; significant after Bonferroni correction). Additionally, paired t-tests comparing border and ectopic variant similarity within each group revealed a significant difference for all but unrelated individuals (p<0.001 each for MZ pairs and DZ/sibling pairs, Bonferroni-corrected). Thus, border variants track more closely with shared genetics than ectopic variants. These findings add to the distinctions seen between border and ectopic variants and suggest that they may have different underlying sources.

**Fig. 6:**
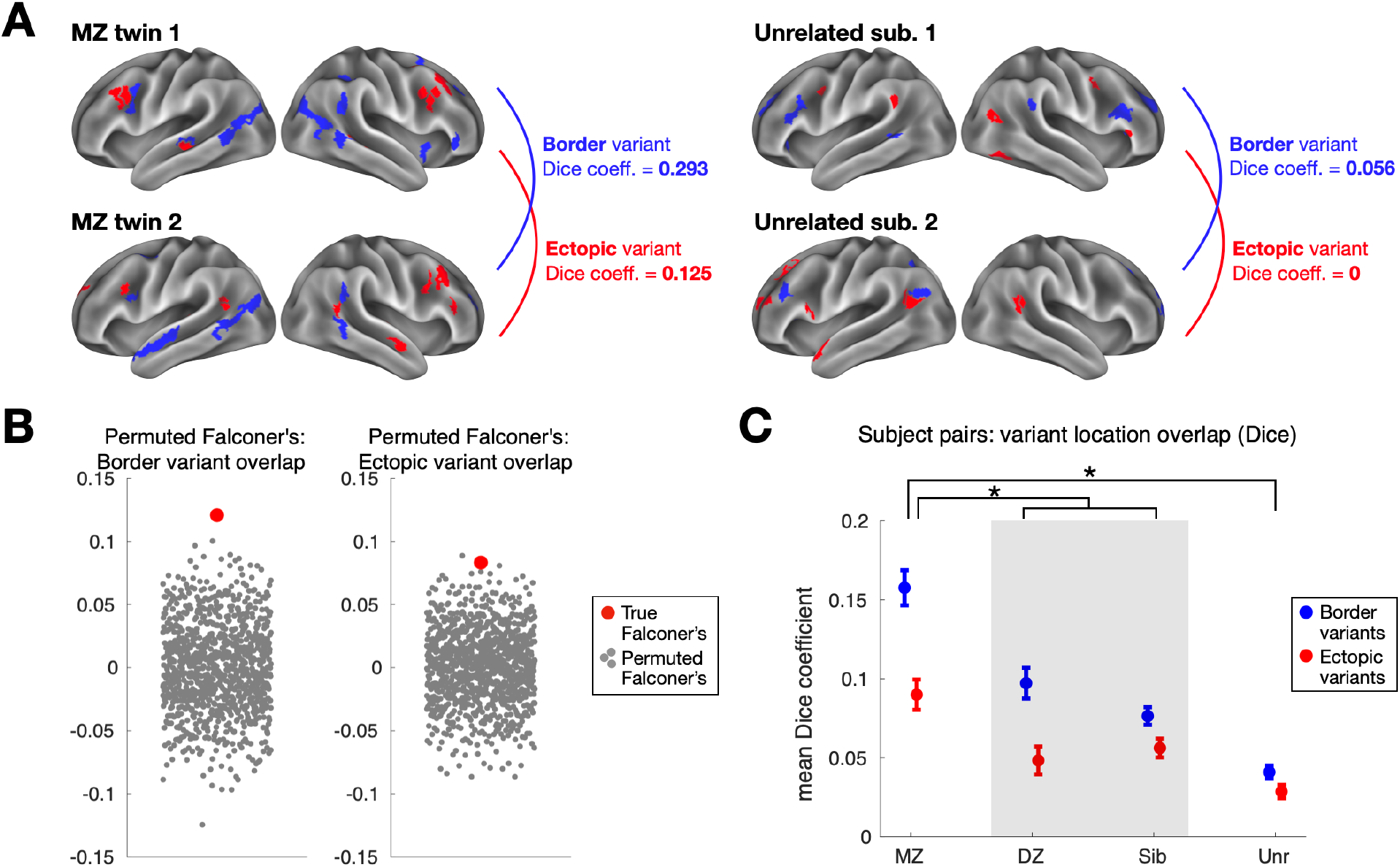
Estimating the genetic influence of border and ectopic variants across individuals in the HCP dataset. (A) The similarity of border (blue) and ectopic (red) variant locations was measured for monozygotic (MZ) and dizygotic (DZ) twin pairs, siblings, and unrelated individuals. This schematic shows an example pair of MZ twins (left) and unrelated individuals (right). The MZ twins exhibit a greater Dice coefficient of overlap between their border variants (in blue) and ectopic variants (in red) than a pair of unrelated subjects. (B) This observation was confirmed by estimating genetic influence with Falconer’s formula, which compares similarity in MZ and DZ twins. Both border and ectopic variants (red dots) exhibited significant similarity (p<0.001 and p<0.002 respectively) relative to a permuted null (gray dots) where MZ and DZ labels were randomly shuffled *(*note, our use of Dice instead of R-values in Falconer’s formula does not produce heritability estimates, but does provide a valid way to assess non-zero genetic influence via permutation testing, randomly shuffling MZ and DZ labels).* (C) Average similarity among border and ectopic variants is shown for pairs of HCP participants. For both forms of variants, MZ twins showed the highest similarity, DZ twins and siblings showed intermediate similarity, and unrelated individuals showed the lowest similarity, a pattern consistent with an influence on genetics on variant locations. Bars represent standard error across subject pairs.

### 2.6 Subgroups diverge between border and ectopic variants

One question that might arise given this evidence for network variants is whether different people share any common properties in their variants. This information may help to constrain theories on how individual differences in brain organization arise across the population. In ref. ^15^, we found that network variants differed across people in a categorical manner, and could be used to identify subgroups of individuals with similar forms of idiosyncratic brain organization. In that work, we found two large reproducible clusters of individuals: those with variants more associated with the DMN and those with variants more associated with control and sensorimotor processing systems. Here, we examined the extent to which similar subgroups would be evident for ectopic and border variants when examined separately. This analysis provides insight into the relative independence between the two variant forms.

To this end, the subset of 374 unrelated individuals in the HCP dataset were grouped based on variants’ similarity to each of 11 network templates and then clustered into subgroups (as in ref. ^15^). This analysis was conducted on two split-halves of the HCP dataset to confirm the reproducibility of data-driven clustering findings. We clustered individuals whose variants had similar network similarity vectors (see *Methods*), identifying three consistent subgroups of individuals in both border and ectopic variants (Fig. 7A and 7B, respectively). The consistency of subgroup assignments for border and ectopic variants (across sessions within an individual) was high (>80% sub-group consistency for both border and ectopic variants; see Supp. Fig. 18), suggesting that the assignments are robust.

**Fig. 7:**
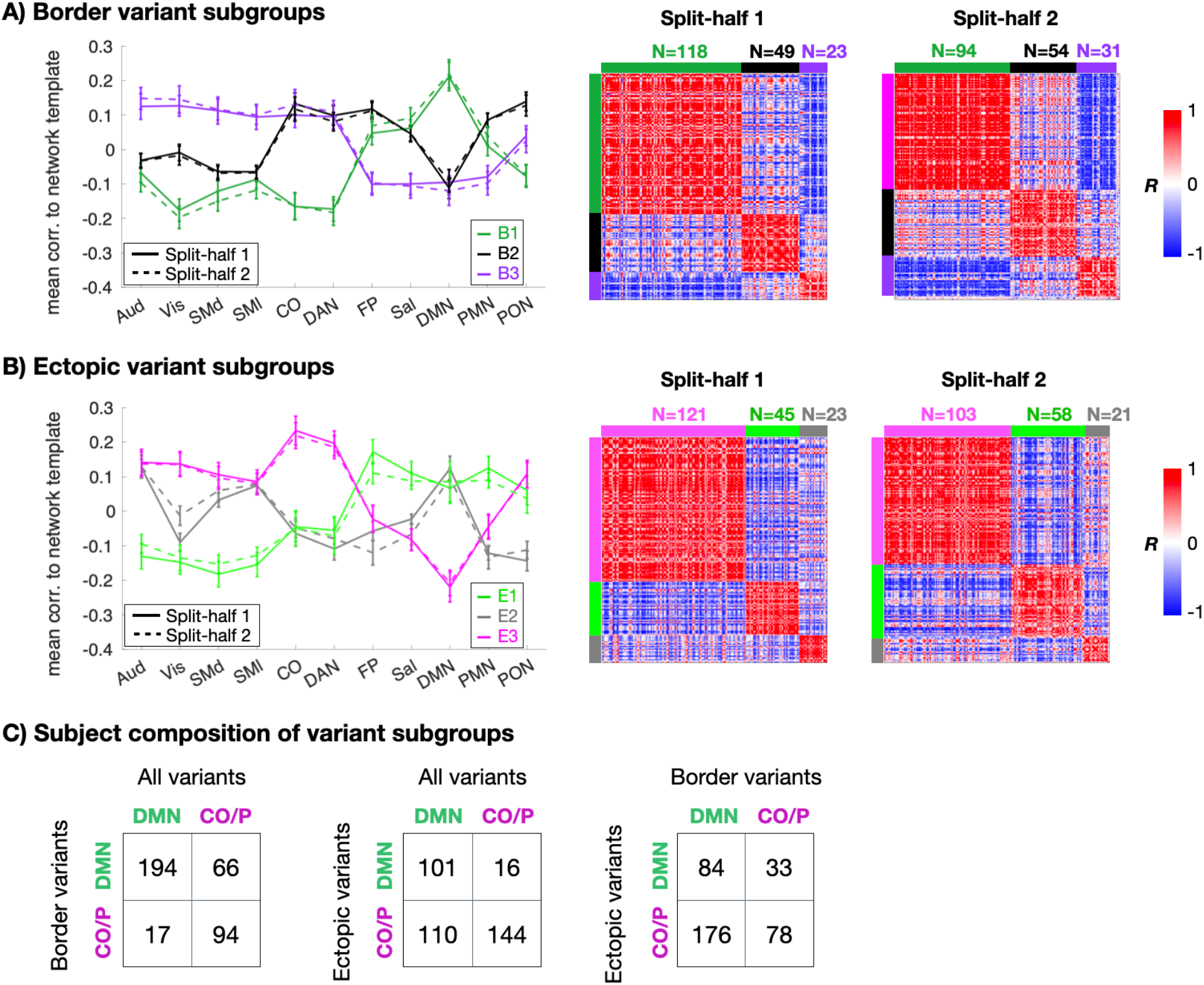
Similarity of border and ectopic variants across subgroups of individuals. For border variants (A) and ectopic variants (B), we separated individuals into subgroups based on the average network similarity vector of their variants (left). Matrices on the right show across-subject similarity (correlation) of variant profiles for each split-half in the HCP. Color blocks at the edges of the matrices denote the subgroup identities. The two variant forms produced three subgroups each with high similarity across matched split-halves of the HCP data. However, the subgroups differed between the two forms. (C) Contingency tables show the composition of subgroups in which each individual’s variant profile (all variants, border variants only, and ectopic variants only) was forced to sort into either a DMN-like subgroup or a control/processing subgroup. Note that ectopic and border variant subgroup labels had poor association with one another.

In clustering individuals via their border variants, we found one large subgroup of individuals whose variants were more highly correlated with the DMN and less highly with control and processing networks (we refer to this subgroup as B1; 57% of subjects, green in Fig. 7A). The second large subgroup had border variants with an intermediate profile, associated with control systems (CO-, DAN-, and FP-like), with a low correlation to sensorimotor networks and the DMN (B2; 28% of subjects, black in Fig. 7A; note this subgroup is distinct from ones observed in our previous analyses). A third smaller subgroup included participants with more CO-like variants, with stronger associations to sensorimotor networks and low correlation to the DMN (B3; 14% of subjects, purple in Fig. 7A; this subgroup was similar to our second subgroup in previous work^15^).

Clustering individuals via ectopic variants resulted in three subgroups as well, but these differed in their specific characteristics. The first subgroup included people with ectopic variants that associated more strongly with FP and DMN and lower correlation to other control networks (E1; 28% of all subjects, light green in Fig. 7B; while similar to B1, note the less prominent DMN profile). A small intermediary subgroup had ectopic variants strongly associated with DMN, auditory and somatomotor networks, and less strongly with control networks (E2; 12% of subjects, gray in Fig. 7B; distinct from any of the border subgroups). The final and largest subgroup had strong associations to the CO, DAN, and PON networks (E3; 60% of subjects, pink in Fig. 7B; most similar to subgroup B3). Thus, our previously published two-subgroup result may have been driven by distinctions between border and ectopic variants.

We next evaluated the consistency of the Infomap subgrouping result relative to two null model tests and relative to the community detection method. When randomizing each subject’s network similarity vector prior to generating the subject-to-subject adjacency matrix, Infomap identified only a single outcome cluster across participants, resulting in significantly higher modularity for the true data relative to 1000 random permutations (p<0.001). When the subject-so-subject adjacency matrix was randomized but preserved strength, degree, and weight distributions (using the *null_model_und_sign* function from the Brain Connectivity Toolbox; www.brain-connectivity-toolbox.net), still the modularity of the true Infomap solution was significantly higher than across Infomap results across 1000 random matrix permutations (p<0.001).

Notably, it appeared that individuals who were members of a particular subgroup in one variant form were not consistently sorted into the same subgroup according to the other variant form; i.e., a subject whose border variants assign them to the DMN-like B1 subgroup was not necessarily assigned to the DMN-like E1 subgroup based on ectopic variants. To confirm this discrepancy, an additional analysis was performed in which each subject was forced into either DMN or control/processing clusters (as these were the most consistently identified across analyses) using a template approach (see *Methods* for more details). All HCP subjects with at least one border and one ectopic variant were included in this analysis (N=371/374). Cluster grouping was then compared for consistency. Figure 7C shows the results of this template-based subgrouping. While an individual’s subgroup based on border variants was somewhat related to its overall (all variants) subgroup (adjusted Rand index = 0.30), subgroups based on ectopic variants were independent of the subgroups based on border (adjusted Rand index = −0.008) or all variants (adjusted Rand index = 0.099). Note that these differences in clustering consistency are substantially larger than seen for a single variant form when compared across sessions (Supp. Fig. 18). This suggests that border and ectopic variant forms are relatively independent, appearing in distinct patterns across individuals.

The results of the subgrouping analysis provide a means of identifying common patterns of variation (or similarity) across individuals. Identifying consistencies of network variants within subgroups of people is relevant for characterizing the basic features of individual differences their differences can inform constraints on individual variation and the forms that they may take in the population. In the same vein, we set out to use the same network-affiliation inputs that were used in this clustering analysis to predict behavioral outputs, asking how the two forms of variation may be able to differentially predict performance.

### 2.7 Border and ectopic variants predict different behavioral phenotypes

In a final exploratory analysis, we examined to what extent border and ectopic variants can predict broad behavioral phenotypes measured outside of the scanner. While these relationships are likely to be small^19, 24, 36, 37^, they may begin to identify subtle differences in the links between these measures of brain organization and measures of cognition and psychopathology.

We used 10-fold cross-validation with support vector regression to predict out-of-scanner HCP behavioral measures (selected to match ref. ^19^) from variant features (see Section 5.11 for extended description). Consistent with past results^19, 24^ we find that it is possible to predict behavioral phenotype from resting-state functional connectivity measures – in this case even in relatively sparse measures restricted to only the most idiosyncratic network locations in each individual (Supp. Fig. 19, Supp. Table 3). The network associations (i.e., the same measures used to sub-group individuals in Section 2.6) of both border and ectopic variants predicted behavioral variables to a small degree (average border prediction r = 0.012, p < 0.04; ectopic prediction r = 0.030, p < 0.001). The strongest predictions were associated with a range of affective, cognitive, and quality of life measures (Supp. Fig. 19A), with ectopic variants showing stronger predictions than border shifts (Supp. Fig. 19B).

Importantly, border and ectopic variants predicted distinct behavioral variables, and no behavioral variables were predicted by both variant forms; predictions between border and ectopic variants correlated at r = −0.126 (Supp. Fig. 19C). Similar differences between border and ectopic variants were seen when the locations of variants were used as features for prediction instead of their network associations (Supp. Fig. 20). Again, no behavioral phenotypes were predicted by both border and ectopic variants (predictions between border and ectopic variants correlated at r = −0.09). Note that, in this case, only border variants were significantly predictive of behavioral phenotypes on average (average r = 0.015, p < 0.007) with the strongest predictions seen for cognitive variables. Notably, similar results were obtained when border vs. ectopic features were defined using the secondary parcellation-free approach (Supp. Fig. 21, 22).

Jointly, these findings suggest that using brain network features to predict behavioral measures is possible, but these relationships are complex and relatively weak on average. The evidence that border and ectopic variants predict different outcomes strongly suggests that simple summary measures (e.g., grouping across all forms of individual differences in brain networks) will mix together distinct brain endophenotypes and muddy interpretations.

As a final test of this last observation, we examined whether joining border and ectopic features together would improve behavioral prediction. Interestingly, despite doubling the number of features in analysis, we found that prediction results were relatively comparable between the joint model and the best-performing individual models (e.g., for network affiliation, ectopic prediction r = 0.030 and joint prediction was r = 0.030). Prediction values were correlated between ectopic and the joint model at r = 0.73. Thus, at least in this case, joining border and ectopic features together does not lead to gains in prediction, but adds ambiguity to the sources of prediction performance.

## 3. DISCUSSION

Individual differences in brain organization have received substantial recent attention. However, most prior studies treat all forms of individual differences as uniform. Here, we document that locations of individual differences (“variants”) come in two forms: shifts in the borders between adjacent systems and ectopic intrusions, islands of an atypical system at a distance from its usual location. Notably, instances of both border and ectopic variants were found in almost all individuals in these datasets – that is, both border and ectopic variants are common properties of brain organization. While they shared some common features, border and ectopic variants differed significantly on a number of properties when contrasted directly, including their spatial location, network assignments, and task-related functional responses. These differences replicated across datasets and approaches to defining border and ectopic variants.

Building on this evidence of their import, both forms of variants showed genetic influences, but border shifts were significantly more similar in identical twins than ectopic variants. Similarly, subgroups appeared independent across the two variant forms. Finally, while both forms of variants could be used to predict behavioral measures collected outside of the scanner, the forms differed in which phenotypes they were linked to. Jointly, these findings suggest that variation in functional brain systems come in (at least) two dissociable forms that appear to link to distinct underlying sources and have differentiable consequences on function. Separation of these two forms is likely to provide new insights into the sources of individual differences and their implications for normative and clinical behavior.

### 3.1. Ectopic islands of variation are common properties of brain organization

A wealth of recent studies has provided evidence of individual differences in brain organization and linked these differences to cognitive variation (e.g., refs. ^19, 25, 38^). The locations of these individual differences are typically treated equivalently, often with the assumption that they reflect differences in the topographic boundaries, or borders, between systems. However, our work suggests that variant brain locations come in two distinct forms – one is associated with border shifts, but another is defined by ectopic intrusions. Ectopic variants are in fact relatively common, comprising ∼40-50% of variants across both datasets. Even at greater distances (e.g., defined at 10mm or further from a similar network boundary), ∼30% of variants were ectopic (see Fig. 2B). Parcellation-free methods also identify robust evidence for ectopic variants that are more than 10mm distant from expected similar locations (Supp. Figs. 3, 6). Thus, not only are ectopic variants common, but many are also observed fairly remote from their expected network boundaries. Many descriptions of individual differences in human brain systems discuss these differences with respect to boundary-related mechanisms (e.g., a region expanding or taking over territory in nearby locations) and distance-based functional alignments have been suggested as a means to address these differences. However, different theories and approaches will be needed to address ectopic variation (see next sections).

Prior to this paper, the existence of ectopic variants had been hinted at in previous work. For instance, ref. ^15^ and ref. ^11^ observed regions where an individual’s functional network organization differs from a group-average description, including regions where a network appeared more distant from its typically observed boundaries (e.g., see DMN variants in Fig. 4B of ref. ^15^ and island of CO appearing in the individual-level map in Fig. 7 of ref. ^11^). Similarly, work by Glasser et al.^39^ parcellating the human cortex noted several instances of sizable spatial displacements in individual-level topography relative to a group-average representation. In this work, we build on these initial observations to systematically characterize the prevalence of ectopic variants and determine how they differ from border shifts along a number of dimensions (e.g., location, similarity in twin samples, and function).

Although common, the proportion of ectopic to border variants varied on a subject-to-subject basis; in the MSC dataset, for instance, some subjects had relatively fewer ectopic variants (i.e., ectopic variants in MSC02 and MSC06 comprised 33% and 40% of total network variants, respectively), whereas in some other subjects (e.g., MSC01, MSC04, and MSC05) the proportion of ectopic variants exceeded 60% (see Supp. Fig. 4 for a breakdown of these proportions in both datasets). Thus, ectopic variants, and more specifically the ratio of ectopic to border variants, may differ systematically across individuals.

### 3.2. Border shifts and ectopic intrusions are distinct forms of individual variation in brain organization

Comparative neuroanatomy studies have shown that cortical functional architecture can differ in a variety of ways across mammals, including differences in the cortical area size/position, number, organization, and connectivity^5^. Although not as often discussed, many of these differences are also seen across individuals within a species^2^. Linking to this work, we hypothesized^15, 40^ that network variants represent a combination of border shifts (expansions, contractions, or displacements relative to the canonical cortical area layout, which will result in differences adjacent to their typical locations) and ectopic intrusions (islands of altered connectivity and function of a region^FN3^ at a distance from the canonical organizational structure). Past work has suggested that even in the typical population, there is substantial variation in the size and position of specific human cortical areas (e.g., in V1 size^22^ with potential links to functional differences in vision^41–43^ and in the position of Broca’s area^44, 45^).

Here we demonstrate that variations both close to and distant from the canonical system structure occur commonly across people and many brain regions, but that these two forms of variation differ along a number of dimensions (in spatial location, network assignment, task activations, genetic influence, prediction of behavioral variables, and subgrouping). These findings were generally consistent across two divergent methods for defining border and ectopic variants, suggesting robustness to the results (see *Supplemental Discussion*). These observations suggest that border and ectopic variants may link to differing underlying sources.

A number of developmental factors influence how cortical functional systems are organized. Gradients in the expression of transcription factors in patterning centers of the cortex control the size and position of many cortical areas^46^. It has been proposed that intrinsic genetic factors create “proto-areas” whose boundaries are refined and sharpened through experience-dependent mechanisms^47^. Similar principles have been theorized to underlie the development of distributed cortical systems, starting from a proto-organization that is refined, fractionated, and sharpened with experience^48^, and groups have reported evidence for heritability in functional brain networks^4, 35, 49–54^. Here we present evidence that the locations of idiosyncratic brain locations are influenced by genetics both for border and ectopic variants, exhibiting higher similarity among identical than fraternal twins. However, variant similarity among identical twins was still far from a perfect identity match, indicating a large contribution of environmental as well as genetic factors to their formation. Interestingly, border shift variants were significantly more similar than ectopic variants in identical twins, suggesting that border shifts may be relatively more linked to genetic factors relative to ectopic variants.

Indeed, while some basic properties are preserved, profound differences in experiences (e.g., sensory deprivation during critical periods) have been shown to substantially alter cortical area size, layout, and connectivity in rodents^5^. Similarly, experience with faces has been demonstrated to be critical to the formation of face selective areas in macaques, although basic retinotopic organization remains^30^. In humans, functional connectivity of congenitally blind individuals shows intact internal topography but large differences in the interareal connectivity of these regions^31^, while those born with only one hand show cortical expansions of regions representing motor functions of other body parts^55^. The lower dependency of ectopic variants on genetic factors suggests that these variants may be more influenced by experience-dependent mechanisms.

These studies in humans and non-human animal models help to explain how cortical organization can both demonstrate substantial commonalities across individuals, but also punctate locations of differences within a species – some associated with local changes (e.g., due to changes in area sizes, which would likely result in border variants), and others that could be linked to more distant alterations (e.g., strong changes in connectivity/function of a (sub)-region that may underlie ectopic variants). Network variants provide a robust and high-throughput approach to identify variations in brain organization across the human brain, helping to constrain theories of the sources and consequences of cortical area variation. Future studies can systematically test the hypotheses raised by this work, using a combination of network modeling methods (for example, testing cascading interaction models for the evolution of network activity^56^), studies of network variants in different age populations (to determine how border and ectopic properties may change over the course of the lifespan), and longitudinal study designs to determine to what extent border and ectopic variants change over time within an individual.

### 3.3. Impact for basic research studies

At present, many resting-state and task-based fMRI studies aggregate or compare data across participants based on spatial normalization, thus assuming that the same spatial layout of brain systems is conserved across individuals^57^. However, widespread individual differences in the localization of brain regions and systems may lead to detrimental effects when performing group-level analyses, including loss of sensitivity and functional resolution^58^, and prediction accuracy of task functional connectivity^59^, task-evoked signals^60, 61^, and prediction of behavior from resting networks^19, 24, 62, 63^. Together, these limitations lead group studies to fall short of providing comprehensive explanations of brain function and cognition as a whole.

Functional alignment across individuals may be improved by more accurate anatomical registration, such as via surface-based mapping methods^64, 65^. However, the resulting functional overlap may be variable depending on the level of cognitive function and brain area in question^23, 66^, with a bias toward enhancing the functional concordance of regions supporting sensory/motor functions^67^. As an alternative, several approaches to increase cross-subject correspondence have been suggested based on improved alignment of functional signals themselves. This includes individualized approaches such as collecting functional localizer task data from each individual subject^57, 68–70^, adopting fcMRI methods to define brain systems and areas from subjects with large quantities of data^12, 71^, or hyperalignment-based techniques to increase functional correspondence^28, 60, 61^. Other methods developed to address functional alignment include template-matching techniques (e.g., refs. ^13, 14, 72^), multi-modal functional/anatomical registration^39^, and hierarchical functional parcellation approaches (e.g., refs. ^19, 24^) to identify brain systems and regions in individuals even with more modest amounts of data.

Each of these methods has demonstrated great utility in allowing us to make cross-subject comparisons to investigate various research questions, including improved definition of default systems^71, 73, 74^, language systems^70, 75, 76^, and sensory-biased frontal regions^77, 78^. However, our work here suggests that they must be implemented in such a way that is able to account for not only local displacements (i.e., proximally altered positions due to border shifts) but also more distant deviations caused by ectopic intrusions. Many current approaches rely on adjusting individual functional regions within a relatively restricted spatial extent, and while this is likely appropriate for many brain locations, imposing a strict distance criterion will not be optimized for detecting ectopic variants that are more distant from their typical network boundaries (see Supp. Fig. 5; nearly one-third of all ectopic variants occur at a distance of more than 30 mm from their same-network boundaries). In particular, our results argue for using procedures that (a) conduct individual-level region identification (e.g., individual localizers, as is often used in the vision science community) or (b) that allow for longer-distance displacements of regions (e.g., with enlarged spotlight procedures). It is also possible that these individual features will link with variable anatomical features (e.g., tertiary sulci^79, 80^) that will provide a new and improved means of alignment of function across people.

Adjusting for ectopic variants will be more relevant in some brain locations and functional networks than others. Compared to border shifts, ectopic variants were much more prevalent in lateral frontal cortex, a region believed to play a role in task control^32, 81, 82^, language^70, 75^, and sensory-biased attention and working memory^77^, among other high-level processes. The prevalence of ectopic variants in lateral frontal cortex suggest that researchers should be particularly cautious in interpreting group-level results in these regions, unless a cross-subject functional alignment method is performed that accounts for non-local deviations. Indeed, important advances in our understanding of lateral frontal cortex will likely be spurred by studies using improved functional alignment methods, which enable separation of multifunctional regions from specialized regions in the face of cross-subject heterogeneity (e.g., refs. ^75, 83^; see review by Smith et al.^84^). In contrast, border variants were more commonly localized to the temporoparietal junction and rostral superior frontal regions, which have been linked to shifting attention^85^ and theory of mind^86–88^ among other functions, suggesting that studies focusing on these domains and areas may benefit from functional alignment approaches which impose distance-constrained changes.

In considering the impact of border and ectopic variants on cognitive neuroscience studies, one intriguing question is the extent to which variant functional connectivity is likely to be changed across task contexts. In our prior work^89^ as well as those of others^90, 91^, we have typically seen that task and rest FC share substantial commonalities, with only relatively subtle changes associated with task states. Similarly, we have demonstrated that network variants are largely stable in their properties across various task states^21^. However, subtle differences in functional connectivity do occur in tasks^89, 92^, and can be used to predict task state^59, 93, 94^, even from individuals^59^. Thus, a fruitful avenue of future investigation will be to establish how tasks affect functional connectivity and border and ectopic variants separately.

### 3.4. Impact for studies of individual differences

One notable difference between the two variant forms is in how they co-vary across participants. In both cases, subgroups of individuals showed similar patterns of variants: for example, with both forms of variants, one subgroup had variants with strong links to the DMN, while another subgroup had variants with stronger links to top-down control networks. However, the subgroups differed in their specific variant profiles (e.g., whether the fronto-parietal network was grouped with the DMN or CO subgroup). Perhaps most notably, the two variant forms appeared independent: that is, a person in the “DMN” subgroup based on border variants could easily be in the “CO” subgroup based on ectopic variants. Other forms of individual differences were also seen in variant properties. For example, there were differences in the relative proportion of each variant form across participants (e.g., MSC02 and MSC06 had relatively few ectopic variants, while MSC01 and MSC05 had many; Fig. 2). These findings beg the question of how each form of individual differences in cortical organization is related to differences in brain function and behavior.

Here, we demonstrated that border and ectopic variant properties are related to robust differences in task activations, not only in the MSC, but also across a range of task activations in the HCP. Border shifts relatively strongly matched the task responses of their associated network, despite their differing location. Ectopic variants, in contrast, showed a more intermediate profile, suggesting that they may be sites of intermediate functional processing.

Consistent with the idea that border and ectopic variants are associated with altered brain function, we also find that these variants can be used to predict behavioral measures collected outside of the scanner in the HCP dataset. While prediction levels were low, they were significant in cross-validated samples. In our previous work, we demonstrated that network variants have high stability across sessions, even up to a year^15^, and across task states^21^. This trait-like characteristic of network variants, as well as its link to differences in functional responses during tasks^15^; Fig. 4) suggest that they are well suited to serve as markers of individual differences in behavior in both the neurotypical population^15^ and in cases of psychiatric and neurological disorders^40^. Other studies have also shown that individual differences in functional brain organization relate to behavior, including links between individual-level network topography and measures of cognition and emotion^19^, associations between changes in functional network topography and a variety of behavioral factors^38^, and links between cognitive ability and maturation of networks supporting executive function^25, 95^.

Intriguingly, we find that border and ectopic variants predict different behavioral phenotypes (with low correlations in their prediction performance). Ectopic variants showed the most robust prediction based on the network associations of variants, and linked to a range of affective and cognitive variables. Border shifts, in contrast, exhibited better prediction based on their locations and were most tightly linked to cognitive measures. Importantly, prediction was not materially improved by joining border and ectopic variant features together. The differences we find between border and ectopic variants suggest that deeper insights into individual differences may be provided if these two forms of variation are separated, or integrated in a more sophisticated manner that acknowledges that they can contribute differing sources of information. This understanding will allow for improved theories about the mechanistic links between individual differences in brain system variation and behavior, likely critical to using this information to guide clinical practice and interventions.

Despite these observations, we note that the brain-behavior predictions reported in this study were relatively small. These results are not surprising given recent findings on the small size of brain-behavior correlations^24, 36, 96^, and our prediction levels are in line with past work that uses similar number/types of features^97^. It may be interesting to speculate on why we find a limited relationship to behavioral measures, and a closer correspondence to differences in task activations. One possibility is limitations in our measurement methods both for our brain measures and behavior^36, 37, 96, 98^. Another is that many individual differences in brain organization are relatively degenerate, producing similar behavioral outcomes (an phenomenon previously termed “behavioral phenocopy”^99^). An important future avenue of research will be to establish principles for how and when individual differences in brain function relate to individual differences in behavioral performance. Regardless of their connection to out-of-scanner behavior, the current findings are important to understand principles of brain organization and how they vary across people. These results will be needed to interpret and form new theories about the neurobiological sources of individual differences.

## 4. CONCLUSION

While the human cortex is organized around a common core architecture, specific locations exhibit prominent deviations from this group-average organization. Here, we investigated two forms of these deviations: nearby shifts in the borders between functional systems and ectopic intrusions at a distance from their typical position. We demonstrate that these two forms of individual variation are both common, but differ in their spatial positions, network assignments, task response patterns, and subgrouping characteristics. Both forms of variants show evidence of genetic influence, but border shifts are significantly more similar in identical twins, suggesting they may be more tightly linked to genetics while ectopic variants may be more influenced by environmental factors. Finally, the two forms of variants predict distinct behavioral variables. These different properties of both forms of variation must be accounted for in the study of cortical system organization and its links to behavior.

## 5. METHODS

### 5.1. Datasets and Overview

Network variants were investigated using data from two separate publicly available datasets: the Midnight Scan Club (MSC) dataset^12^, and a subset of individuals from the Human Connectome Project (HCP)^100^. The MSC dataset is a “precision” fMRI dataset consisting of 10 highly sampled subjects (5 female; average age 29.3 years; 1 participant excluded due to high motion and sleep^12^) with over 154 minutes of low-motion rest fMRI data and task fMRI data across 3 conditions (mixed, memory, motor – in this manuscript we focus on the results of the mixed design tasks). From the larger HCP dataset, we primarily analyzed 384 unrelated subjects (210 female; average age 28.4 years as in ref. ^15^, selected to be unrelated and have a minimum of 45 min. of low motion resting-state fMRI; see SI Table 1 in ref. ^15^ for details on exclusion criteria for this dataset). Of these 384 subjects, 374 were retained for analysis after removing subjects with exceptionally low spatial correspondence to the group-average indicating low quality data (see Section 5.3; note however that results remain the same prior to their removal). For analysis of behavioral links and genetic influence on variant locations, a larger subset of the full HCP dataset was used. This group consisted of 823 individual subjects (793 after quality control), including those with familial relationships, each with a minimum of 40 min. of low-motion resting-state fMRI data.

In compliance with ethical regulations, informed consent was obtained from all participants. Study protocol for the MSC dataset was approved by the Washington University School of Medicine Human Studies Committee and Institutional Review Board, and protocol for the HCP dataset was approved by the Washington University Institutional Review Board.

In both datasets, previously defined idiosyncratic locations of functional connectivity (“network variants”)^15^ were divided into homogeneous segments (see criteria below) and segregated into ectopic intrusions (“ectopic variants”) and border shifts (“border variants”) based on the criteria described below. Several features of each variant form were then examined. First, we quantified the prevalence of both variant forms across the two datasets. Second, we characterized the spatial location and idiosyncratic (individual-specific) network assignment of these regions. Third, we examined the task responses of both variant forms using task data from the MSC dataset. Fourth, we examined if there were common profiles of border shift and ectopic variants (as we have seen in past data for variants as a whole^15^), by clustering individuals into subgroups based on their variant characteristics.

### 5.2. Preprocessing

Imaging data from the MSC and HCP subjects used in the present analyses were preprocessed identically to ref. ^15^. Full details on acquisition parameters, preprocessing, FC processing, and volume-to-surface mapping can be found in that manuscript, but are outlined briefly below.

Functional data from both datasets were preprocessed to remove noise and artifacts, following ref. ^101^. For the HCP dataset, we began with the dataset as processed following the minimal preprocessing pipelines^102^. Procedures included field map distortion correction of the functional images, slice-timing correction (for the MSC dataset only), mode-1000 normalization, motion correction via a rigid body transformation, affine registration of functional data to a T1-weighted image, and affine alignment into stereotactic atlas space (MNI for HCP (Montreal Neurological Institute, Montreal, QC, Canada); Talairach for MSC^103^).

Following this, resting-state fMRI data was further denoised for functional connectivity analysis, including regression of white matter, cerebrospinal fluid, and whole brain signals, six rigid-body parameters and their derivatives, and their expansion terms^104^. High-motion frames (calculated as framewise displacement; FD^105^) were censored; frames with FD > 0.2 were censored for the MSC data^12^ and frames with filtered FD > 0.1 were censored from the HCP data following ref. to address respiration contamination of motion parameters (filtered FD = low-pass filtering at <0.1 Hz of the original motion parameters prior to FD calculation; note that two participants in the MSC dataset – MSC03 and MSC10 – with strong respiratory contamination of their motion parameters also used the filtered FD measure^12, 89^). As in ref. ^107^, 5 frames at the start of each run along with any segments < 5 frames long were also removed. These censored frames were interpolated over using a power-spectral matched interpolation. Subsequent to this, a temporal bandpass filter was applied to the data from 0.009 to 0.08 Hz.

Following this processing, BOLD data were mapped to each subject’s native cortical surface as generated by FreeSurfer from the atlas-registered T1^108^. Data were registered into fs_LR space^66^ and were aligned to the surface following Gordon et al.^7^, producing a CIFTI file with a BOLD timeseries for each functional run. From this point on, all analyses were conducted on the cortical surface (at the vertex-level).

Data on the cortical surface was spatially smoothed with the application of a geodesic smoothing kernel (σ = 2.55; FWHM = 6 mm). Finally, high motion frames (that were previously interpolated) were removed from analysis. Note that participants were required to have at least 40 min. of data total to be retained in analysis (see refs. ^15, 21^ for evidence that ∼40 minutes of data is necessary to achieve reliable network variant measures). Some additional improved reliability is seen beyond 40 min., however. This factor motivated our use and replication of results in both the HCP and MSC datasets (when possible), in order to balance the advantages of large numbers of participants with the added reliability of extended amounts of data.

### 5.3. Defining network variants

Network variants are defined as locations in an individual that show strong differences in their functional network patterns relative to the group average. In the current manuscript we began with the same set of variants as originally presented in ref. ^15^. Briefly, for each subject, each vertex’s seedmap (i.e., its cortex-wide connectivity map) was correlated with the same vertex’s seedmap^FN4^ from a group-average reference dataset, composed of an independent 120 young adults (the WashU 120^14^). After repeating this procedure for all cortical vertices, this produced one individual:group similarity map per person (with a spatial correlation value for each vertex). HCP subjects whose average individual:group spatial correlation value was greater than 2 standard deviations below the mean (r = 0.436) were excluded from further analyses (10 out of the primary subset of 384 subjects; 30 out of the larger pool of 823 subjects used for analyses of similarity within twin samples and behavior).

Similarity maps were thresholded to include only the lowest decile of correlation values (i.e., to identify the 10% of locations where the individual was least similar from the group-average) and were then binarized. The decile criterion was originally selected because it represents the approximate inflection point on a histogram of individual-to-group similarity, suggesting that it may serve as a natural criterion for identifying points with strong differences from the group average. In past work, we have tested other methods for selecting network variants (based on absolute rather than relative criteria or different relative thresholds^15, 21^), with similar overall results. As in ref. ^15^, a vertex-wise map of low-signal regions was used to mask out potential variant locations. Clusters of at least 50 neighboring vertices in size were flagged as pre-variants for further analysis (see Fig. 1A).

Following this original variant definition, a series of steps was taken to further refine the set of pre-variants to generate the final variants that were used in the border/ectopic analyses, given observations that some pre-variants were large and irregularly shaped, suggesting they might consist of separate units. To divide pre-variants, previously defined contiguous units were divided into segments with the goal of minimizing heterogeneity in the connectivity of individual pre-variants. This procedure consisted of a two-fold check of (1) the variance explained by the first principal component of the pre-variant, resulting from a principal component analysis on variants’ vertex-wise seed maps (i.e., ‘homogeneity’ ^7^) and (2) the proportion of the variant’s territory that is dominated by a single network in the individual’s subject-specific vertex-wise network map. In these vertex-wise maps, each cortical vertex is individually assigned to a network using a template-matching procedure (see refs. ^13, 14, 72^ which matches each vertex’s thresholded seedmap to each network’s thresholded seedmap (each thresholded at the top 5% of values) and assigns the vertex to the network with the best fit (measured via the Dice coefficient; similar to ref. ^13^).

Using the MSC as a pilot dataset, two independent raters visually assessed all subjects’ variants by examining connectivity seedmaps of vertices within a pre-variant. Based on the apparent homogeneity or inhomogeneity of seedmaps at various locations within the pre-variant, the raters evaluated whether each pre-variant should be flagged to be divided. Based on these results, an algorithm was developed that best matched the hand ratings from reviewers, based on a combination of template-match network representation and homogeneity thresholds. The thresholds were then applied to the broader set of MSC and HCP participants. These final thresholds were set at 66.7% homogeneity and 75% network dominance in the individual network map. The thresholds were then applied to the broader set of MSC and HCP participants. Flagged pre-variants were split along the network boundaries of the vertex-wise network, resulting in final “split” – but each contiguous – variants into homogeneous regions. Clusters smaller than 30 contiguous vertices were removed. See Fig. 1A for a schematic representation of the splitting procedure and examples of split variants. Following the combination of variant definition, size and SNR exclusion, and pre-variant homogeneity refinement, approximately 2% of vertices were defined as variants in each individual.

### 5.4. Functional network assignment of variants

Variants were then assigned to a best-fitting canonical functional network by a procedure which matched each variant to its best-fitting functional network template as in ref. ^15^. To assign variants to networks in the MSC dataset we used group-average network templates that were generated in previous work from 14 networks using data from the WashU 120 (refs. ^13, 15^; see Supp. Fig. 12A). Networks used for MSC analyses included default mode (DMN), visual, fronto-parietal (FP), dorsal attention (DAN), language (Lang.; note this has been referred to as the ventral attention network in our past work but we have now reclassified as language based on its correspondence with language localizers^76^), salience, cingulo-opercular (CO), somatomotor dorsal (SMd), somatomotor lateral (SMl), auditory, temporal pole (Tpole), medial temporal lobe (MTL), parietal medial (PMN), and parieto-occipital (PON). For terminology, variants whose best-fitting functional network is DMN based on comparisons to all network templates, for example, are termed “DMN variants.”

For the HCP dataset (given differences in dataset resolution and acquisition parameters^100^; see also ref. ^72^), a dataset-specific network template was generated (Supp. Fig. 12B; see ref. ^15^ for template generation procedure). The networks included in HCP-specific analyses were similar to those for the MSC, but did not include the language, MTL, or T-pole networks as these networks did not emerge consistently across edge density thresholds from the data-driven group-average network identification procedure (Infomap^109^). In both cases, the average seedmap for each variant in an individual was compared with each of the network templates (after binarizing both to the top 5% of connectivity values (as in ref. ^13^) and assigned to the template with the highest Dice coefficient overlap. Network variants were removed from further analysis if they did not match to any functional network (Dice coefficient of zero) or if over 50% of their vertices overlapped with the group-average network for that location.

### 5.5. Classification of variants as ectopic intrusions or border shifts

Two methods were used to classify variants as either ectopic intrusions or border shifts. The primary method was implemented to identify variants which lay adjacent or at a distance from the canonical group-average regions of the same network. First, using the MSC as a pilot dataset, all variants were manually classified as ectopic variants by examining whether they visually appeared to be spatial extensions of existing network features or whether they appeared to arise unconnected from other same-network locations. Next, we tested different geodesic distance criteria to optimize agreement of computed border/ectopic classifications with manual classifications of variants as border/ectopic based on our visual inspection of the MSC participant variants. The final procedure specified that all variants further than 3.5 mm (edge-to-edge distance) away from a same-network cluster were classified as ectopic variants; variants closer than 3.5 mm were classified as border variants. This procedure was then applied to the independent HCP dataset (see Fig. 1 for schematic representation). As a check to determine whether the 3.5 mm distance threshold we specified would significantly impact the proportion of ectopic variants in our sample, we also classified variants as ectopic or border based on a distance criterion of 5, 7.5, and 10 mm and computed the proportion of ectopic variants at each. Finally, we also quantified the distance between each final ectopic variant and canonical regions of their assigned network.

A secondary, parcellation-free method to classify variants was also implemented for use in a subset of analyses (reported in the supplement). Rather than relying on a group parcellation to make a border-ectopic distinction, this technique classified a variant based on whether its average connectivity seedmap reached high similarity to the group average seedmap (i.e., high variant-to-group *R*) at brain locations near the variant (a border shift), or whether the variant’s similarity to the group did not approach a peak in correlation at any brain location near the variant (an ectopic intrusion). This procedure specified that if the seedmap of a variant reached at least 90% of its peak correlation to the group average at a distance within 10 mm (edge-to-edge distance), then the variant was classified as a border shift. Conversely, a variant whose seedmap similarity to the group did not approach at least 90% of its peak correlation within a distance of 10 mm was classified as ectopic. In both cases, the “peak” variant-to-group *R* was defined as the most similar seedmap comparison within 150 mm of the variant. For a schematic illustration of this method, see Supp. Fig. 2.

Of note, each of these variant classification methods relies on a comparison between an individual and a group-average description (a group-average network map in the primary method, or a group-average connectivity map in the secondary method). The motivation for this approach was twofold: first, we sought to understand how group-level representations may err, both in terms of proximal adjustments (border shifts, which can be more straightforwardly addressed by available techniques) and for more distant, individually idiosyncratic locations (ectopic intrusions, which arise further away than expected based on group priors). Second, if we are to approach the question from a theoretical standpoint, we may assume that an individual may deviate from a “standard” template of brain organization in various ways. Some mechanisms may result in relatively proximal border shifts (e.g., differences in the transcription factor gradients which mediate a standard set of developmental pathways), whereas others may operate over longer distances (e.g., experience-dependent competitive mechanisms). Thus, it is important to characterize each of these possible routes for individual variation. However, to contextualize an individual’s variants within their *own* individually defined network map, we provide an exploration of variants in the MSC dataset with ectopic variants classified into 3 sub-types based on how isolated or connected the variants were to their individual-specific networks; see Supp. Fig. 7 and legend for an illustration, and Supp. Table 1 for classification results.

### 5.6. Examining differences in spatial distribution between ectopic and border variants

To visualize the spatial distribution of border and ectopic network variants, a spatial overlap map was generated by summing network variant maps (separated by form) across individuals. This produced an overlap map highlighting regions of high and low occurrence of variants in both the primary and parcellation-free classification method.

These spatial overlap maps were quantitatively contrasted at two levels. First, an omnibus map-wise permutation analysis was run. Our null hypothesis was that the distribution of border and ectopic variants was no more different than would be expected by chance. We used a permutation approach to address this hypothesis. This was achieved by (1) shuffling variant classification labels (ectopic vs. border) randomly at the subject level (i.e., flipping variant labels within a subject’s variants map 50% of the time) to create pseudo-ectopic and pseudo-border variants, (2) summing variant locations across subjects to generate a cross-subject overlap map for pseudo-ectopic variants and a cross-subject overlap map for pseudo-border variants, (3) calculating the similarity (spatial correlation) between the pseudo-ectopic and pseudo-border overlap maps, and (4) repeating steps 1-3 1000 times for different permuted labels. We then compared the distribution of permuted similarity values with the true similarity between ectopic and border spatial distribution maps. We calculated significance as a *p-*value based on the proportion of permutations in which the permuted correlation value exceeded the true correlation value. This procedure was used to determine whether spatial distributions, as a whole, differed between ectopic intrusions and border shifts. Permuting border and ectopic labels at the subject level allowed us to preserve spatial relationships and local spatial auto-correlation^110^ within each subject’s variant map, in turn allowing us to directly test the null hypothesis that the distributions of border versus ectopic variants across subjects are no more different than would be expected by chance.

Second, to locate specific regions where the distribution of ectopic variants was significantly distinct from border variants, a cluster-size-based permutation analysis was run. The first two steps were the same as before: (1) shuffling ectopic and border labels within each subject’s variant map to create pseudo-ectopic and pseudo-border variants in the same proportion and (2) summing variant locations across subjects to create an overlap map for pseudo-ectopic and pseudo-border variants for each of 1000 permutations. Using these permuted overlap maps, we (3) generated 1000 difference maps of the pseudo-ectopic variants distribution minus the pseudo-border variants distribution, (4) thresholded these pseudo-difference maps to only keep locations with differences of at least 5% of participants (19 subjects), and (5) calculated the size (number of vertices) of retained clusters. This procedure produced 1000 permuted pseudo cluster size calculations. This distribution of cluster sizes was used to define a cluster threshold that corresponded to the top 5% of permuted (random-chance) clusters (p < 0.05 cluster-corrected). Finally, the true ectopic–border difference map was also thresholded to only keep locations with a difference of at least 5% of participants, and all clusters composed of fewer vertices than the cluster-correction threshold (229 vertices) were removed, and the resulting cluster-corrected difference map is displayed.

### 5.7. Comparing network assignments between variant forms

In addition to its spatial location, each variant has an idiosyncratic functional network assignment (see “Functional network assignment of variants” section above; e.g., a network variant located in a typical DMN region may have functional connectivity more closely associated with FP, causing it to be assigned to that system). In the next analysis, we used a permutation approach to quantify differences in border and ectopic variants’ functional network assignments.

For this analysis: (1) we permuted the label of each variant as border or ectopic within participants in the HCP dataset (permutations were done per participant), (2) we then calculated the proportion of pseudo-ectopic to pseudo-border variants for each network, (3) repeated steps 1-2 1000 times. We then compared the true ectopic to border proportion for each network to the permuted distribution of proportions. Significance was assessed via *p-*values calculated as the proportion of permutations in which a network’s true percentage of ectopic variants was less or greater than (two-tailed) all permuted percentages after FDR correcting for multiple comparisons across networks. This analysis was also performed using the parcellation-free variant labels. To further test whether frequencies of each variant form varied by network, a mixed-effects generalized linear model analysis was performed. Factor effects were defined for variant form (border or ectopic) and variant network, with one level per subject recording counts of each observed combination. The interaction model was performed to measure the extent to which the influence of variant form on variant frequencies varied across networks, while accounting for within-subject dependence.

Finally, we determined the “swaps” of network territory occupied by border and ectopic variants (e.g., a variant located in canonical cingulo-opercular territory that “swaps” its network assignment to the fronto-parietal network). To do so, we defined each variant’s consensus network assignment as the modal network across variant vertices in the pre-defined group-average system map, compared this with the variant’s assigned network, and tabulated the frequency of all cross-network swaps.

### 5.8. Examining task activation of variants

In addition to defining FC features of network variants, we also examined how these regions responded during tasks. Following Seitzman et al.^15^, we first focused on fMRI task activations during the mixed-design tasks in the MSC dataset (semantic and coherence), given strong a-priori hypotheses about the responses of different networks during these tasks. The semantic task involved participants indicating whether presented words were nouns or verbs, and the coherence task required participants to indicate whether an array of white dots on a black background were displayed in a concentric (as opposed to random) arrangement. Within each task block, a short cue signaled the onset of the block, with a series of individual trials presented with jittered timing. Another short cue signaled the end of the block, and task blocks were separated by fixation periods of 44 seconds (see ref. ^12^ for more details on task design).

Task fMRI data from the MSC dataset underwent the same basic preprocessing as listed in the *Preprocessing* section (i.e., field map correction, slice timing correction, motion correction, alignment, and normalization, registration to the cortical surface and smoothing). These tasks were then analyzed using a general linear model (GLM). For each event (cues, correct and error trials of each type), eight separate timepoints were modeled in a finite impulse response modeling approach^101^); for each task block, a block regressor was modeled to estimate sustained activations. The tasks and analysis streams are described in further detail by refs. ^12^ and ^89^.

In this study, we examined the activation image across all conditions (start/end cues, trials, and sustained activations across semantic and coherence tasks) versus baseline to interrogate network variant locations to examine whether forms of variants exhibited differences in task activations. A series of comparisons were conducted following ref. ^15^: (1) comparisons (two-tailed t-tests) of the task activation of DMN variant locations in a given subject relative to the same location in other subjects, (2) a comparison of task activations of variant locations in each network relative to canonical regions of their assigned network, and (3) a comparison of task activations of variant locations in each network relative to canonical regions of *other* networks. Task activations were examined separately for ectopic variants and border variants; this analysis was repeated using the parcellation-free variant labels.

Task fMRI data from the HCP dataset were also used to query the activation properties of border and ectopic variant regions. We used the MSM-Sulc registered, 4mm-smoothing versions of task contrast images from the publicly available analysis-level data from all 7 tasks (emotional processing, gambling, language, motor, relational processing, social cognition, and working memory), originally processed following the HCP preprocessing pipeline (ref. ^102^; see section 5.2). All tasks used a blocked design to model each contrast; see ref. ^112^ for details. Of the primary set of 374 subjects, 358 who had both forms of variants and data available for all tasks were retained for the task activation analyses. Again, we compared task activations of variant locations in each network relative to both canonical regions of their assigned network and canonical regions of *other* networks (this time separately per contrast and per network, given the expanded number of contrasts). Networks with fewer than 200 variants overall were excluded to increase the stability of the results.

To determine the extent to which each form of variant exhibited shifted task activations relative to the typical response expected for canonical regions of their assigned network, we calculated the proportion of this shift (normalized by the average activation seen across all other networks) as follows:

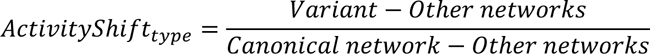

In order to assess the variants’ *relative* shift, contrasts were only included in this analysis if they showed a difference in the given network’s activation compared to all other networks’ activation; only contrasts with a difference of at least 0.5% signal change were considered. This analysis was performed in both datasets: in the MSC summarized across networks, and in the HCP summarized across contrasts within a network.

### 5.9. Similarity of network variants in twin samples

We next asked about potential sources for these different forms of idiosyncratic variation in functional networks. We used the familial design of the HCP data to interrogate this question. Thus, unlike in previous analyses, we included all N = 793 participants who passed our quality control criteria (see sections 5.2 and 5.3), not excluding those with familial relationships. Of these, 784 subjects with at least one border variant and at least one ectopic variant were carried forward in the twin analyses (*note: only MZ and DZ twin pairs whose twin status was confirmed via genotyping were included as twins*).

Next, we estimated the variant similarity among twin samples by contrasting the Dice overlap in monozygotic (MZ) and dizygotic (DZ) twins using Falconer’s formula for broad sense heritability^34^: h^2^ = 2(D_MZ_ – D_Dz_). The formula is derived from the model that MZ twins share 100% of their genes and DZ twins share 50%, while both share common environmental variables. Note that our input values to Falconer’s formula were Dice overlap coefficients and not r values, and as such they cannot be directly contrasted with Falconer’s estimates based on correlation, but still provide a valid means to test non-zero genetic influence. To assess the significance of our estimate of genetic influence, we used a non-parametric permutation approach, in which MZ and DZ labels were randomly shuffled before Falconer’s formula was re-computed. This was done 1000 times to create a null distribution. P-values were then calculated relative to this null distribution. These analyses were repeated using variant labels from the parcellation-free variant classification method.

Finally, we then compared the locations of border and ectopic variants across individuals with different familial relationships (monozygotic twins, dizygotic twins, siblings, and unrelated individuals) using the Dice overlap coefficient. A two-way mixed effects ANOVA was performed to test for an interaction between variant form and group, with groups associated with 100% genetic overlap (MZ twins), 50% genetic overlap (DZ and non-twin siblings; note that for this analysis, Dice values from DZ and non-twin sibling pairs were combined due to approximately equivalent genetic relatedness and the smaller N of DZ pairs alone), and 0% genetic overlap (unrelated individuals). Following the ANOVA, two-sample *t*-tests were performed, assuming unequal variance, to assess between-group differences of border minus ectopic Dice coefficients (e.g., MZ_border–ectopic_ vs. combined DZ/sibling _border–ectopic_ vs. unrelated _border–ectopic_). Bonferroni correction was applied to these three tests. Finally, a paired t-test was performed to test whether, in each group, there was a difference between border and ectopic variant similarity among pairs. Again, Bonferroni correction was applied to these three tests.

### 5.10. Identifying subgroups of ectopic and border variants across individuals

We next sought to investigate whether there were commonalities across subgroups of individuals in their network variant characteristics. To this end, we conducted a similar analysis as in ref. ^15^ to examine the subgrouping potential of ectopic and border variants. In this case, data was restricted to the sample of N = 374 unrelated individuals to avoid confounding subgrouping analyses with potential familial relationships examined in section 5.9.

As in ref. ^15^, we split the HCP dataset into 2 matched samples consisting of 192 subjects each (further pared down to 183 and 191 subjects, as described in Section 5.3), allowing us to search for data-driven findings that replicate across split-halves. In every individual in the HCP dataset, the mean similarity between all variant vertices and each network was produced by calculating the correlation between the variants’ seedmap and each of 11 network template seedmaps. This created an 11 x 1 vector for each subject containing information on the network similarity of variant locations. These vectors were correlated between all subjects in each split half, producing a subject adjacency matrix. A data-driven approach, the Infomap clustering algorithm^109^, was then conducted on the cross-subject correlation matrix in one split-half; the algorithm was applied after thresholding the correlation matrix across a wide range of density thresholds (5% to 50%, in increments of 1%, requiring a minimum subgroup size of 20 subjects). Subgroups were defined based on consistent results across a range of Infomap thresholds. Sub-group assignments were validated in the second split-half of subjects by correlating each subject’s network vector with one of the resulting subgroup average network vectors; a minimum correlation of 0.3 was required for assignment. In this analysis, we opted to use Infomap as our method of community detection as it has been shown to out-perform other methods – including modularity-maximization approaches – on benchmark testing for community detection^113^. Infomap better addresses issues that arise from resolution limits, with an improved ability to identify modules of different sizes.

After running Infomap, three primary subgroups were consistently produced across a broad range of thresholds; also note that additional subgroups can be identified at sparser thresholds). Subgroups of individuals with similar network similarity profiles were identified in the first (discovery) split-half, and the subgroups were replicated in the second (validation) split-half as described above. Variant network connectivity patterns of resulting subgroups of individuals were subsequently examined. The subgrouping analysis was performed twofold: first operating solely on ectopic variants, then operating solely on border variants.

Next, we examined to what extent an individual’s subgroup assignment was consistent when grouped by all of their variants, their border variants only, and their ectopic variants only. All HCP subjects with at least one of each form of network variant were included in this analysis. The network similarity vectors of two previously identified stable subgroups (a DMN subgroup and a control/processing subgroup) were identified by clustering individuals’ network vectors across all variant forms within each split-half, and averaging across split-halves. The DMN and control/processing network similarity vectors were then used as templates with which each individual’s network variant profiles were correlated (separately across all variants, border variants only, and ectopic variants only). Similar network similarity results were found in original analyses based on all variants^15^; see Supp. Fig. 22 for DMN and control/processing profiles. Following these “forced” subgroupings, the adjusted Rand index was calculated three-fold to investigate any similarity between an individual’s subgroup assignment between (1) all variants and border variants only, (2) all variants and ectopic variants only, and (3) border variants and ectopic variants.

We next investigated the robustness of an individual’s subgroup assignment, asking whether this assignment was consistent within a subject when their resting-state data was divided in two. For this analysis, a participant’s data was split into two parts, one for each session, and an 11 x 1 network-similarity vector containing information on border and ectopic variant locations (as described above) was produced for each session’s data. The data-driven subgroup profiles (network-similarity vectors, each sub-group profile averaged across split-halves as in Fig. 7A/B) were correlated with each subject’s network-similarity vector (separately for border and ectopic variants). For each subject, the subgroup profile with the highest correlation was designated the match for a given session’s data, and a contingency table was produced to illustrate the proportion of subjects whose split-session data yielded an equivalent subgroup assignment (see Supp. Fig. 18).

Finally, we sought to evaluate the quality of our Infomap clustering in two ways. First, we randomized the 1x11 vector of network associations for each subject, and used the randomized subject-to-subject correlation matrix to cluster individuals. This process was repeated 1000 times to produce a null distribution of 1000 permuted modularity values of the clustering solution, which were compared to the modularity of the true clustering solution. Second, we used the *null_model_und_sign* function from the Brain Connectivity Toolbox (www.brain-connectivity-toolbox.net) to create a subject-to-subject adjacency matrix matched in degree, weight, and strength distributions to the original input matrix (using the default of 5 swaps of each edge); this was again repeated 1000 times and compared to the true modularity.

### 5.11. Predicting behavioral phenotypes from network variants

An additional set of analyses modeled after ref. ^19^ were conducted to determine how border and ectopic variants predicted behavioral measures collected outside of the scanner. For this set of analyses we used the full set of HCP participants that passed our low-motion threshold and quality control criteria (N = 784, the same subset used for analyses described in section 5.9), including all twin and non-twin siblings, in order to maximize our sample size given recent evidence that cross-sectional brain-behavior associations require large samples to be robust^36^.

Our primary analyses focused on using the network affiliations of variants (as in section 5.10) to predict 58 behavioral variables in the HCP as in ref. ^19^; see Supp. Table 3 for a full list of variables. Prediction features from this analysis were based on the average affiliation of variants to 11 template networks. That is, for each person, we estimated the extent to which their variants (on average) were correlated with the 11 template networks, producing 11 continuous feature values that ranged from –1 to 1. This same affiliation measure was used to sub-group individuals in the analyses described in 5.10, connecting with that work. Separate prediction analyses were conducted for each form of variants (border and ectopic), and for both forms together. Supplemental analyses also examined how the location of border and ectopic variants (vectorized binary map) predicted behavioral performance.

We used 10-fold cross-validation for prediction, accounting for familial status in the creation of folds (i.e., related individuals – based on either father or mother ID -- were kept in the same fold to ensure independence across folds). Within each fold, we used support vector regression in Matlab (*fitrsvm*) with default parameters to identify a relationship between brain features and behavioral variables; this relationship was then tested on the left-out test fold. As in ref. ^19^, we regressed out age, sex, BMI, mean FD, and DVARS from the behavioral measures prior to prediction as these measures correlate with scanner motion^114^ (regression was carried out independently in the training data and then applied to the test data to prevent leakage across folds^19^). Prediction accuracy is reported as the correlation between the predicted and true behavioral measures in independent test data. Correlations were measured per fold and averaged over folds.

We used permutation testing to assess the significance of predictions. For each permutation, behavioral variables were randomly reordered across subjects, thereby breaking the link between variant measures and behavioral outcomes in each participant. Then, the same support vector regression with 10-fold cross validation approach was carried out, resulting in the average correlation between true and predicted values in permuted data. Permutations were repeated 1000 times (500 times for supplemental data based on variant locations) to generate a null distribution that was used as a benchmark for the predictions in the true data. These analyses were repeated using variant labels from the parcellation-free variant classification method.

## 6. Data availability

Data from the Midnight Scan Club is publicly available at https://openneuro.org/datasets/ds000224. Imaging data from the Human Connectome Project (1200 Subjects Release) can be accessed at https://db.humanconnectome.org/; some data elements utilized in this work (e.g., family structure, behavioral measures) require second-tier permissions from the HCP for access. Data associated with the WashU-120 is available at https://openneuro.org/datasets/ds000243/versions/00001.

## 7. Code availability

Code for original network variant definition and primary analyses is available at https://github.com/MidnightScanClub. Code used to analyze border and ectopic variants will be made available upon publication at https://github.com/GrattonLab.

## Acknowledgements

Funding was provided by NIH grant R01MH118370 (CG), the James S. McDonnell Foundation (SEP), NIH grant R01MH111640 (MN), NIH grant T32MH100019 (ANN), the Therapeutic Cognitive Neuroscience Fund (DMS), and NIH grant K01AA030083 (ASH).

## Footnotes

FN1 Network variant locations are present even after surface-based normalization^63, 64, 101^ that align data across people by large-scale sulcal features. Individual differences in fcMRI are not well related to variations in anatomical metrics (refs. ^7, 15^; although it is possible that they relate to finer-scale anatomical features, e.g., see ref. ^100^). This dissociation from gross anatomical features, together with correspondence to task responses, suggests that variants may relate more closely to differences in the positions of functional brain areas or systems, which can vary relative to anatomical landmarks^23, 65^.

FN2 A note on terminology: variants of a given network assignment are henceforth referred to as, e.g., “DMN variants,” referring to the variant’s network association (that is, locations that are not in typical DMN regions but have a seedmap that matches canonical DMN distributions). Regions that are, instead, found in canonical regions of the DMN are referred to in that way without any shortening.

FN3 Note that while we have made efforts to identify homogeneous network variants, each network variant is not necessarily equivalent to a full cortical area. Consider the example of border shift variants: these variants could arise because an area within an individual has been expanded or displaced across a border to subsume territory of another region. Using our methods, only the non-overlapping segment will be labeled a variant. Therefore, even if functional connectivity were a perfect proxy for brain area divisions, these variants might represent only a subunit within the area.

FN4 Note that for these analyses we focused on cortex-wide seedmaps, not separating edges from within and between network connections. In our experience, focusing on whole brain maps and top within-network connections produces similar results^72^. It will be interesting in future work to more fully explore the differences between defining variants on each form of edge.

