## Supplementary Material for "Two common and distinct forms of variation in human functional brain networks"

### SUPPLEMENTARY MATERIALS

#### SUPPLEMENTAL DISCUSSION

##### *Comparison between methods for defining border and ectopic variants*

While our original identification of variant locations is defined without reference to a hard parcellation (i.e., based on the continuous similarity to the group average connectivity map at that location), our primary method of then separating variants into border and ectopic sub-types is done with reference to the distance from a (hard-parcellated) canonical network map. Given this dependency, and that multiple distinct canonical network maps exist (e.g., (Power et al., 2011; Yeo et al., 2011)), we developed a secondary method for defining network variants that did not depend on a particular networks parcellation (see Supp. Fig. 2).

For the most part, the results across these methods were quite consistent. Both methods identified a sizeable proportion of variants as ectopic (Fig. 2, Supp. Fig. 3). Both methods identified very similar spatial distributions of border and ectopic variants (Fig. 3, Supp. Fig 10), with more ectopic variants in the right lateral frontal cortex and more border shift variants near the temporo-parietal junction and in superior frontal cortex). Both methods also find similar shifts in the task activation profiles for border and ectopic variants (Fig. 5, Supp. Fig. 16), with border variants more strongly shifted than ectopic variants. Both methods also show a similar pattern of heritability across border and ectopic variants (Fig. 6, Supp. Fig. 17) and similar behavioral prediction results (Supp. Fig. 19-22).

Some differences between the methods were present as well. These include slight differences in the proportion of variants defined as border and ectopic (e.g., compare Fig. 2A to Supp. Fig. 3). This is likely driven, at least partly, by the greater distance criteria we used in the parcellation free method, as the proportions found in this method are comparable to the 10 mm. distance results from our primary method (e.g., compare to Fig. 2B). Furthermore, while both methods identified similar associations for variants (Fig. 4, Supp. Fig. 14A) with most variants assigned to the DMN, FP, and CO networks, the ratio of border-to-ectopic variants per network varied somewhat across methods (Fig. 4B, Supp. Fig. 14B). Thus, future work examining this metric should consider including multiple methods for defining network variants to establish stability of the result.

In general, each approach that we used has advantages and disadvantages. Parcellation-dependent approaches tend to be easier to implement and to interpret, leading to clearer understanding of what border and ectopic variants represent. However, these approaches are dependent on the quality and resolution of the original parcellation. While the parcellation that we use has been extensively validated in both group averages (Power *et al.*, 2011) and individuals (Gordon et al., 2017b; Laumann et al., 2015), it makes specific choices regarding the resolution of networks to be examined (e.g., (Gordon et al., 2020)). In contrast, parcellation-free approaches such as the one that we implemented here can be implemented without selecting a particular network definition and resolution. However, these approaches still require the selection of thresholds for defining border and ectopic variants (e.g., 90% peak similarity at 10 mm.), are computationally intensive to implement, and can sometimes be more challenging to interpret (e.g., there may be multiple reasons for differences in peak similarity architecture). In practice, a combination of both approaches may prove useful in establishing robust properties of border and ectopic variants, as highlighted with the results from the network-affiliation analysis.

**SUPPLEMENTAL FIGURES**

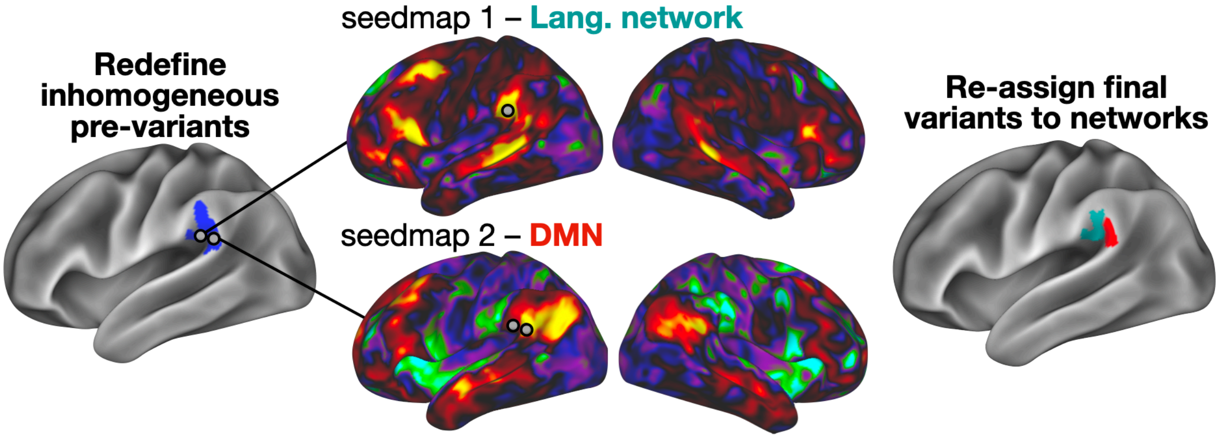


*Supp. Fig. 1: Refining of inhomogeneous pre-variants.* Pre-variants were flagged to be divided if either of two criteria indicated high heterogeneity (a PCA "homogeneity" measure (Gordon et al., 2016)) and a measure of network territory in the subject's network map - see *Methods*). For example, the pre-variant in this panel included two sub-regions, one with relatively high affiliation to the DMN and one with high affiliation to the language network. Flagged pre-variants were then divided along their network map sub-divisions, resulting in a set of final variants for each subject.

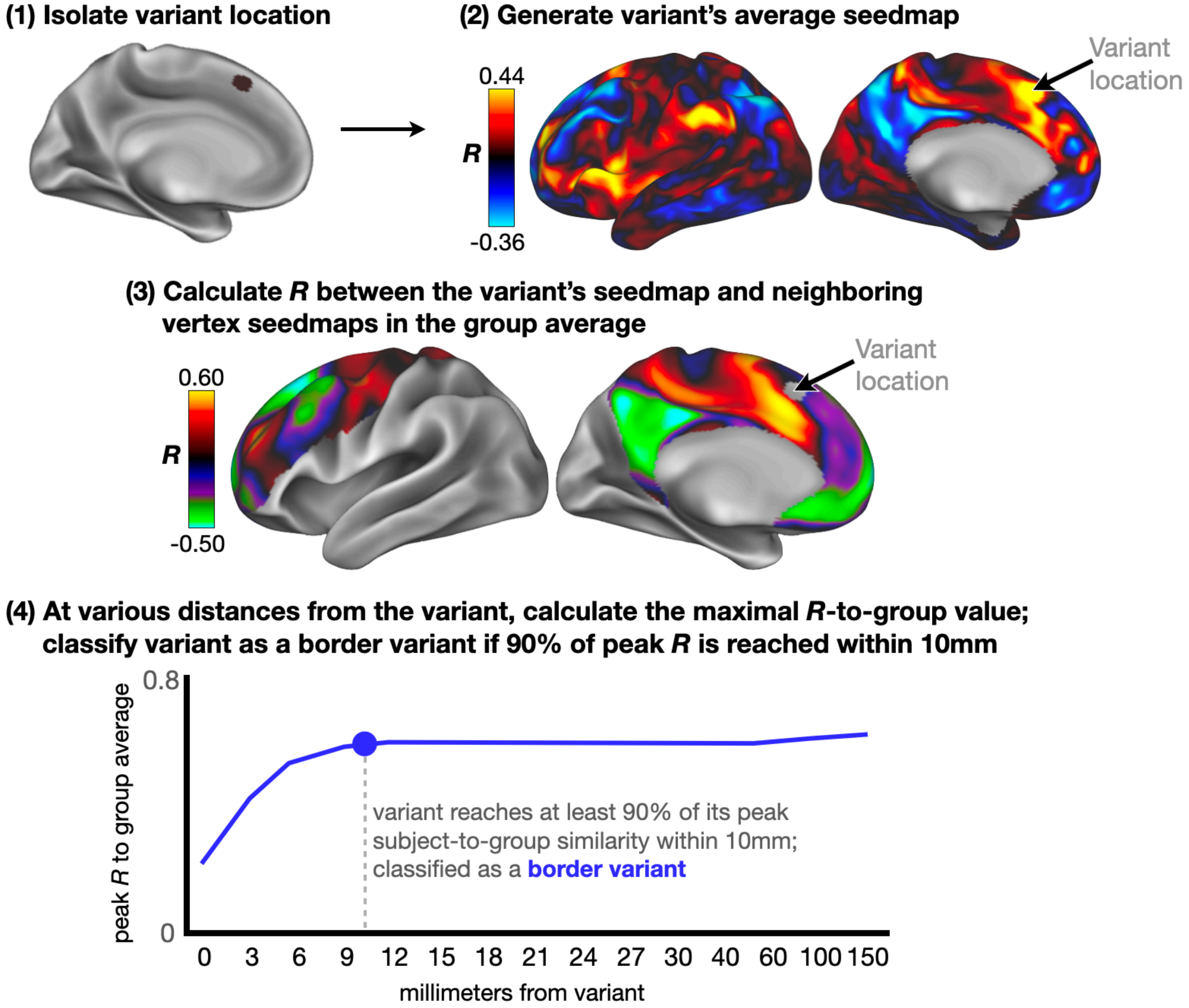


*Supp. Fig. 2: Secondary method for classifying border and ectopic variants.* To classify variants as border or ectopic without relying on a specific group parcellation of functional networks, a parcellation-free method was implemented. First, (1) a variant's location was isolated using the methods described in Fig. 1A. (2) We then generated the average seedmap for the variant (averaged across vertices within the variant). (3) The variant's seedmap was compared via spatial correlation to group-average seedmaps of brain locations within 150 mm (edge-to-edge distance) from the variant. The schematic shows similarity between each displayed vertex (in the group average) and the variant location; in this image the variant location in question appears to show highest similarity to nearby locations within the group-average (i.e., the nearby dACC). (4) The similarity was quantified across a range of distances up to 150 mm; at each distance the maximum variant-to-group *R* value was calculated. A variant was classified as a border shift if it achieved at least 90% peak similarity to the group by 10 mm, and as an ectopic intrusion if 90% of peak similarity was not reached by 10 mm. Peak similarity was defined as the maximal correlation between the variant seedmap and the (group-average) seedmap of any vertex located within a distance of 150 mm.  Note that this approach does not require a pre-set definition of canonical networks (e.g., default mode, frontoparietal, etc.), bypassing differences between common group parcellations (e.g., Yeo (Yeo *et al.*, 2011) vs. Gordon (Gordon et al., 2017a)) and resolutions (Yeo 7 vs. 17 (Yeo *et al.*, 2011)).

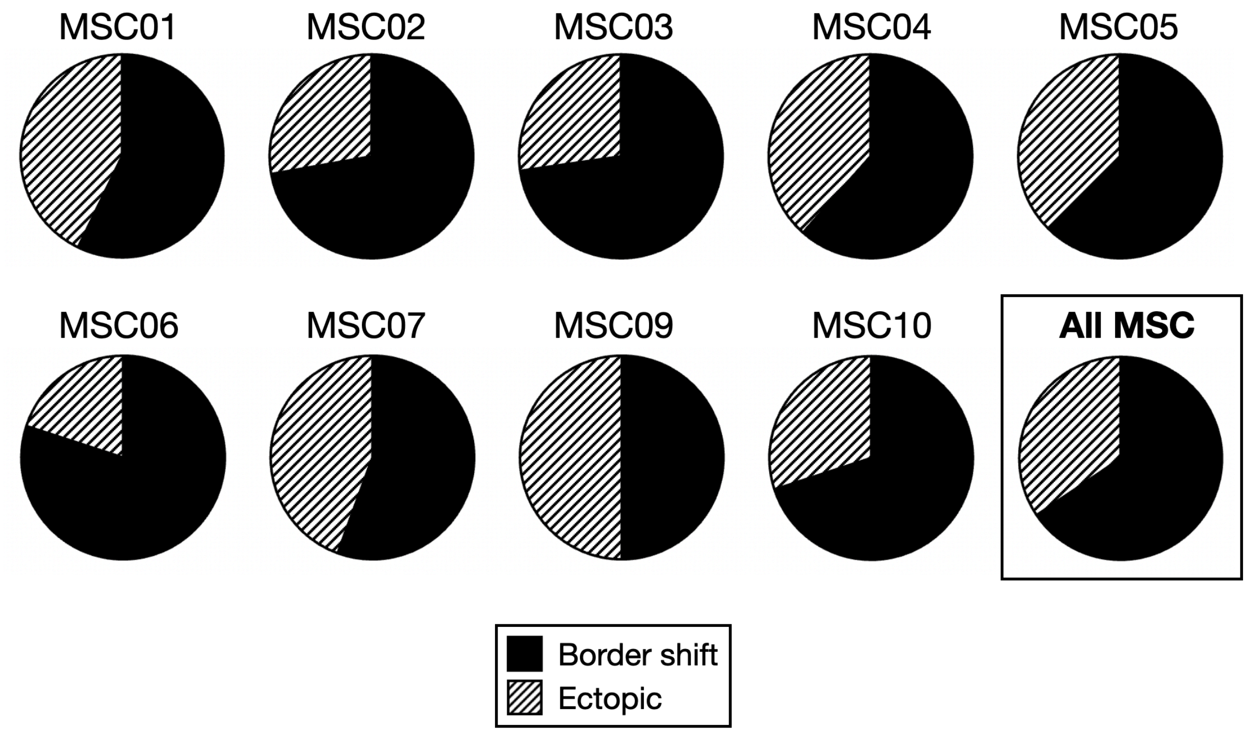


*Supp. Fig. 3: Proportion of border and ectopic variants in the MSC dataset using the parcellation-free classification method.* As in the primary method (Fig. 1A), we identified examples of both border-shift and ectopic variants in each MSC participant. The parcellation-free method was slightly more conservative in identifying ectopic variants than the primary method, probably due to the larger distance criteria (i.e., both this method and the primary method at 10 mm. identified roughly a third of variants as ectopic).

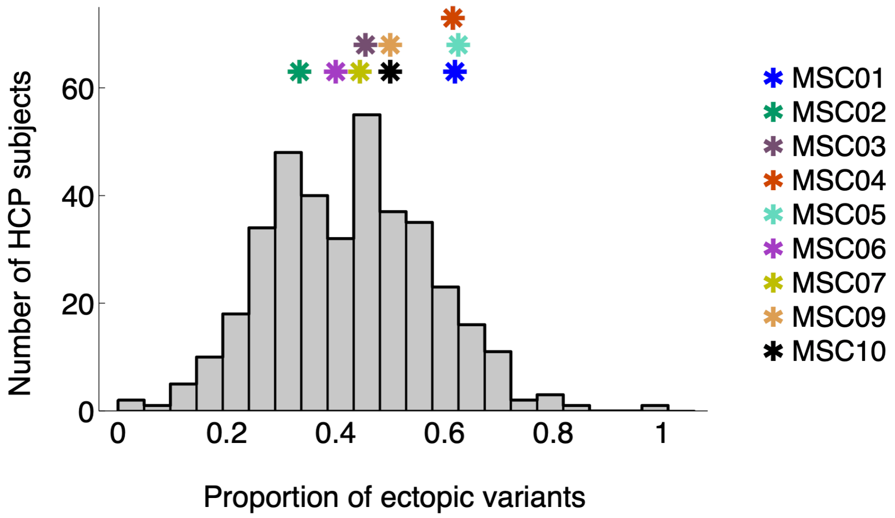


*Supp. Fig. 4: Proportion of ectopic variants across subjects.* The histogram shows the proportion of ectopic variants (out of total variants) across all subjects in the HCP dataset using the primary definition approach from the manuscript. Stars on top show the proportion of ectopic variants for each subject in the MSC dataset (labeled at right).

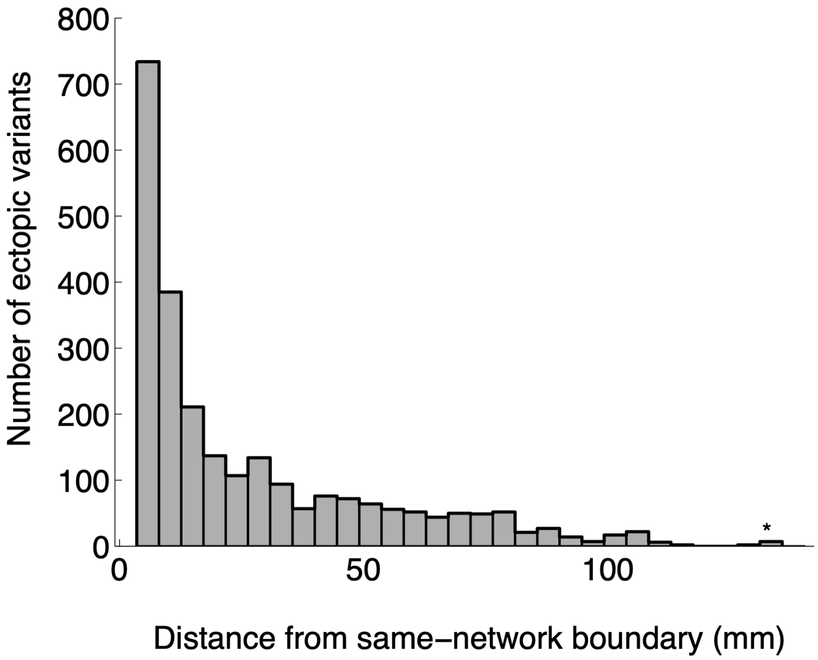


*Supp. Fig. 5: Histogram of ectopic variants' distances from their nearest same-network boundary.* The plot depicts a histogram of the edge-to-edge distance between each variant and canonical regions of a given network in the HCP dataset using the primary border/ectopic definition approach from the manuscript. A sizable proportion of ectopic variants are found at a long cortical distance for consensus locations of their network (median = 15.5 mm; N = 2499 variants), consistent with the results of the secondary definition method (Supp. Fig. 6).

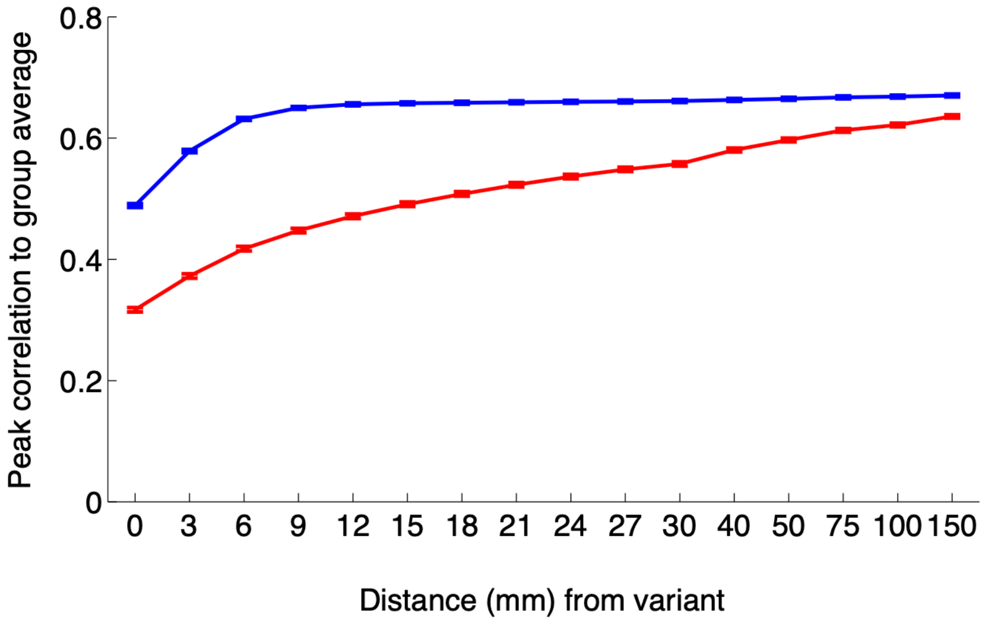


*Supp. Fig. 6: Parcellation-free variant definition: similarity-to-group curves.* Implementing a parcellation-free method to classify border and ectopic variants reveals that ectopic variants are less similar to the group-average at locations nearer to the variant, and only achieve comparable similarity to the group at higher distances. Variant similarity data were averaged across border or ectopic variants within a subject, then averaged across all 374 subjects; error bars represent standard error across subjects.

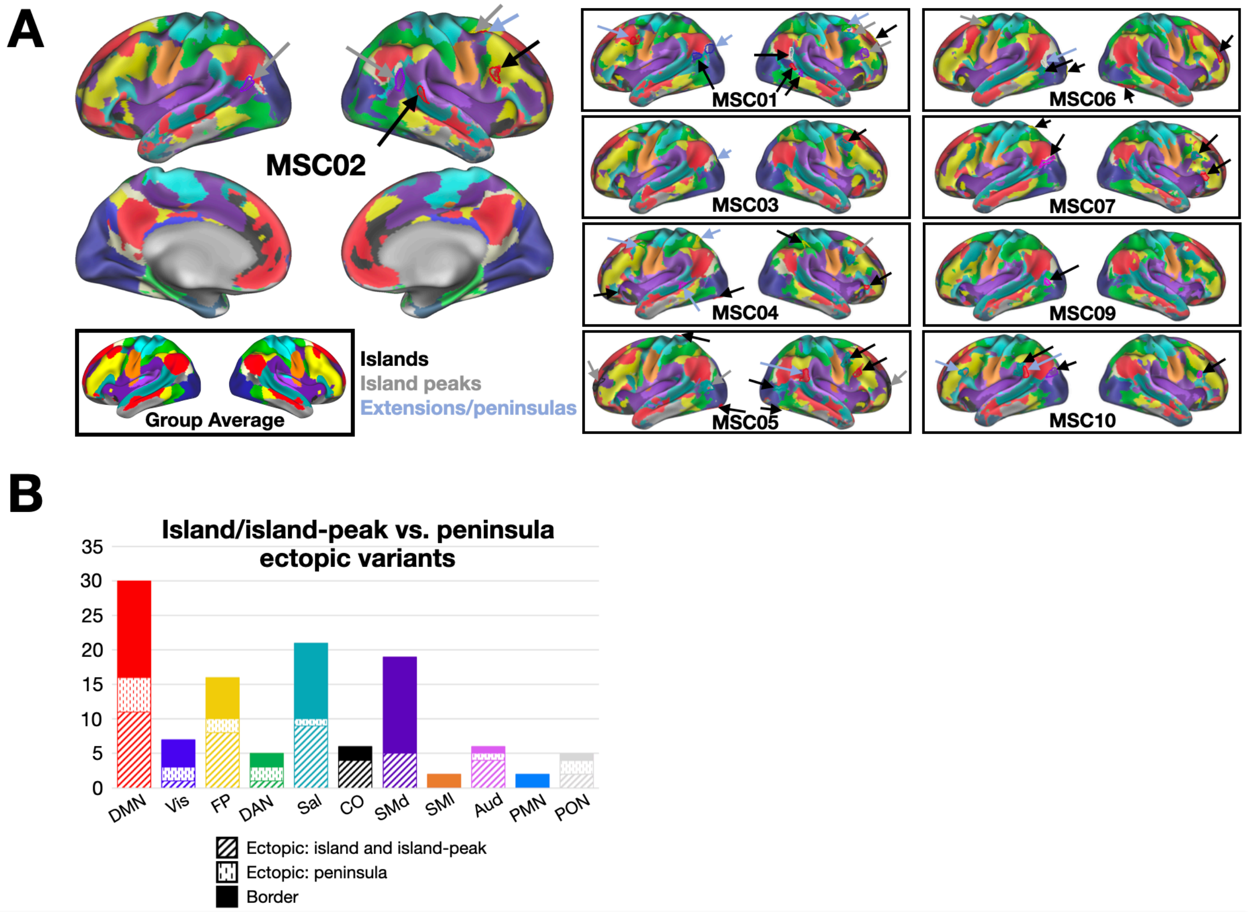


*Supp. Fig. 7: Exploration of ectopic variant sub-types in the MSC, relative to individually defined networks*. (A) Variants are shown as outlines, with individual network maps shown in filled in colors in the background. Each variant was classified as an island (black arrows; regions isolated from major segments of their own network), island peak (gray arrows; regions isolated from major segments of their own network, but with the variant representing <50% of the region), and extensions/peninsulas (blue arrows; regions connected to larger swaths of their own network). Most ectopic variants were either islands or island peaks in the MSC (Supp. Table 1). (B) Network assignments of ectopic variant sub-types across the MSC. While a portion of the ectopic variants in the MSC dataset were classified as peninsulas, the majority remained islands and island peaks.

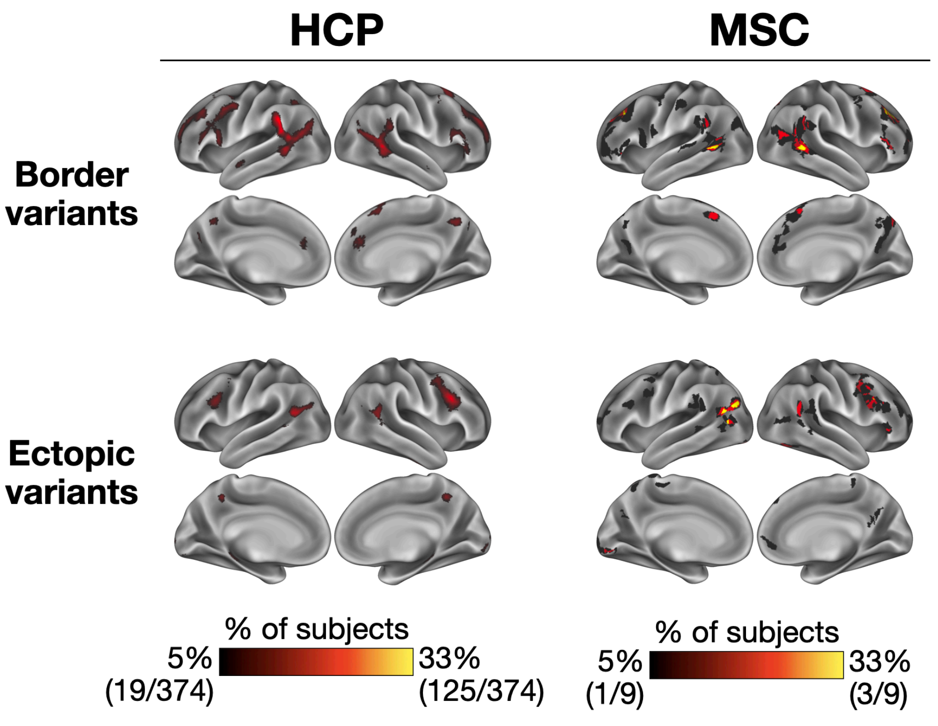


*Supp. Fig. 8: Side-by-side comparison of border and ectopic variant spatial distributions between the HCP and MSC datasets*. Both forms of variants appear in many of the same regions, including ectopic variants in lateral frontal regions in the right hemisphere and border variants around the temporoparietal junction and superior rostral frontal regions in both hemispheres.

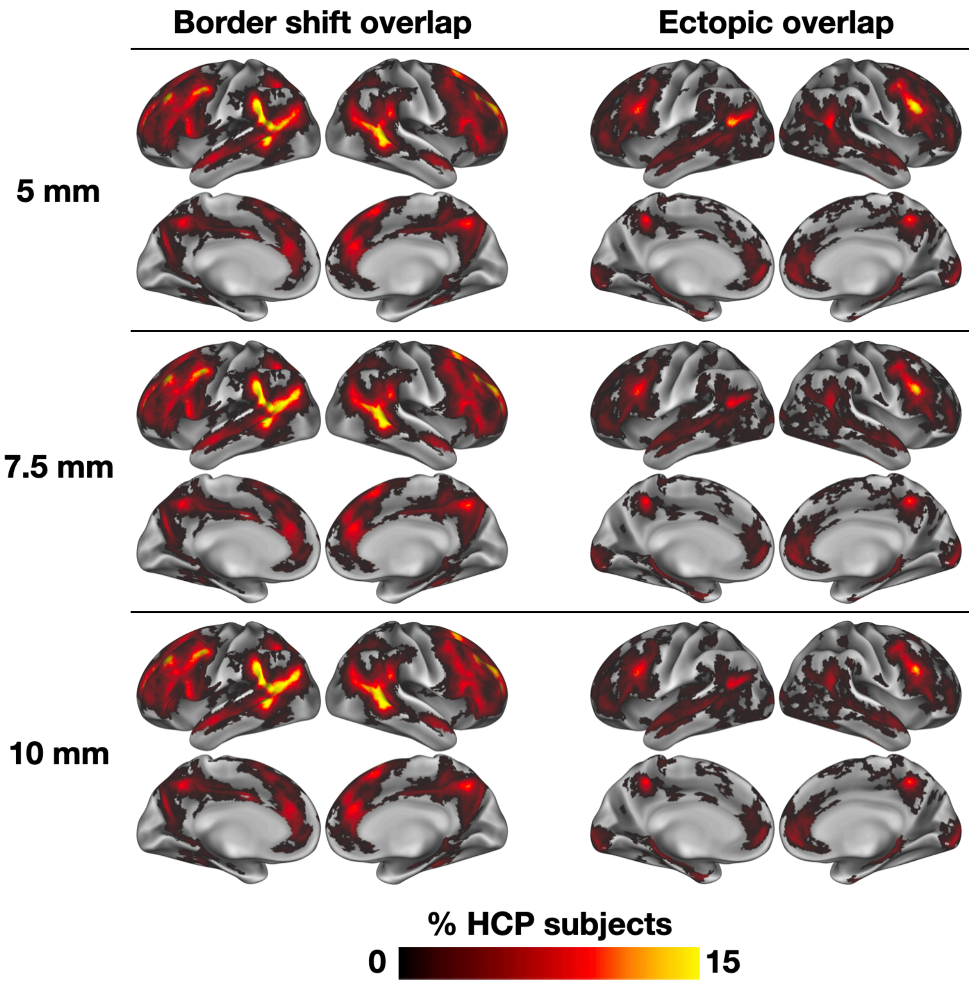


*Supp. Fig. 9: Properties of ectopic variants as a function of increasing distance requirement for classification.* When using the primary approach to define ectopic variants, the spatial distribution patterns observed when ectopic variants in the HCP dataset are defined at > 3.5 mm from network borders (i.e., Fig. 3A in the main manuscript) are largely conserved even as ectopic variant distance is increased through 10 mm.

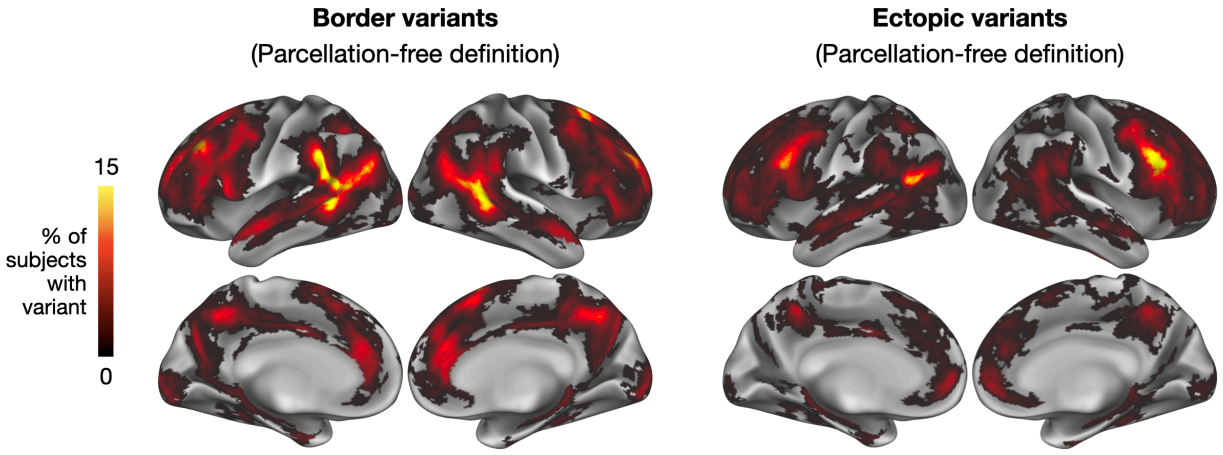


*Supp. Fig. 10: Spatial distribution of ectopic variants in the HCP dataset using the parcellation-free classification method.* When the parcellation-independent method for classifying border and ectopic variants is implemented, similar spatial distribution patterns are observed relative to the primary classification method.

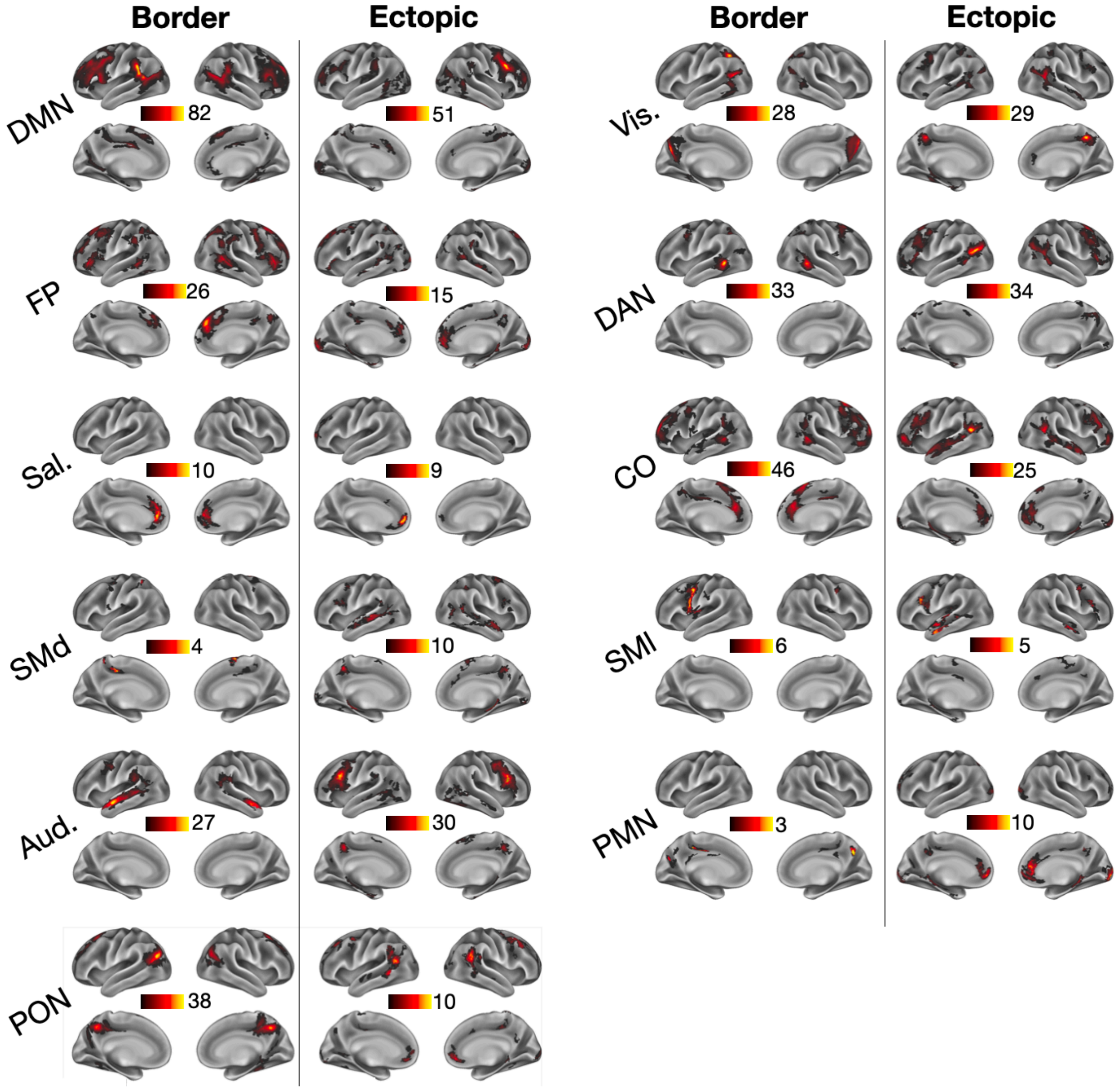


*Supp. Fig. 11: Border and ectopic variant spatial overlap maps across HCP subjects, separately for each functional network.* Note that the color scales differ for each network's plot (maximal subject overlap values for each network are displayed), given different baseline rates associated with variants in each network.

A)

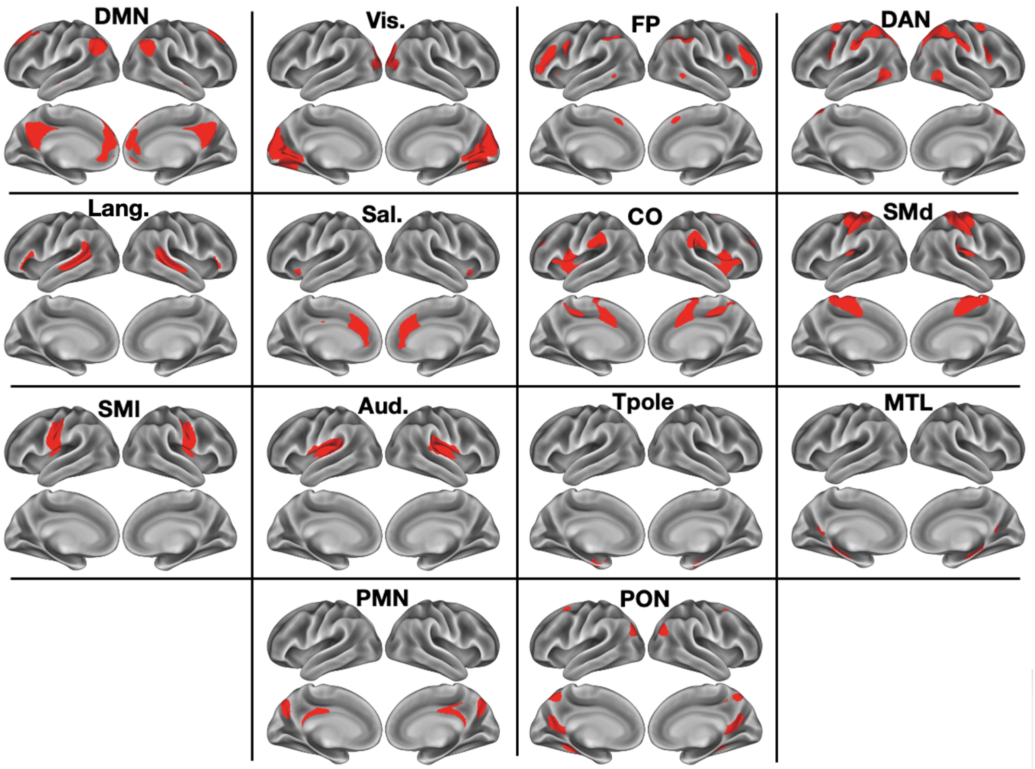


B)

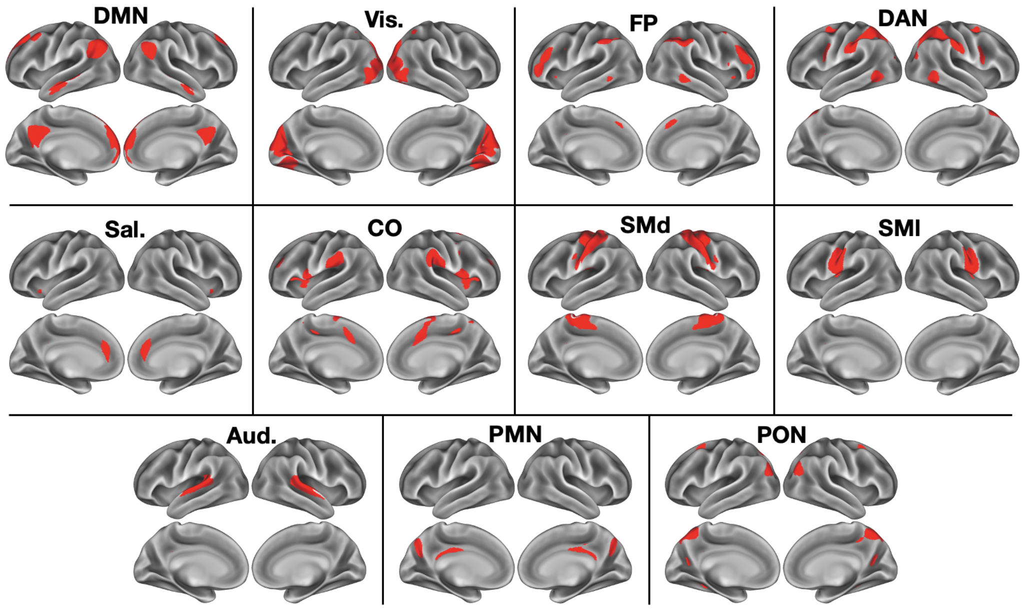


*Supp. Fig. 12: Network templates.* (A) Network templates derived from the WashU-120 dataset (used for MSC analyses). (B) Network templates derived from the HCP dataset (used for HCP analyses). See *Methods* and (Seitzman et al., 2019) for further details on the generation of network templates.

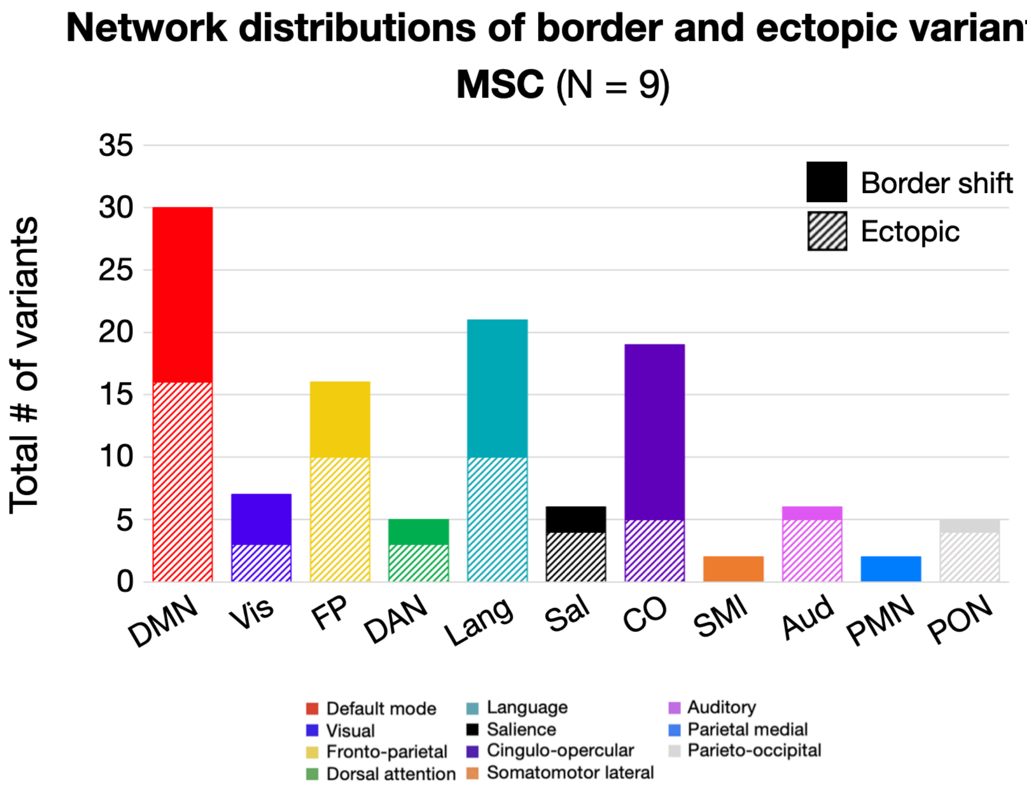


*Supp. Fig. 13: Network distributions of border and ectopic variants in the MSC dataset.* Each bar shows the number of variants associated with 12 common association networks in the MSC dataset, separated by whether variants were border shift or ectopic (similar to Fig. 4 in the main text). Note that in the MSC, many variants are also associated with the Language network, but this network template was not identified in the HCP dataset (see *Methods, Supp. Fig. 12).* The primary border/ectopic definition approach was used for these analyses.

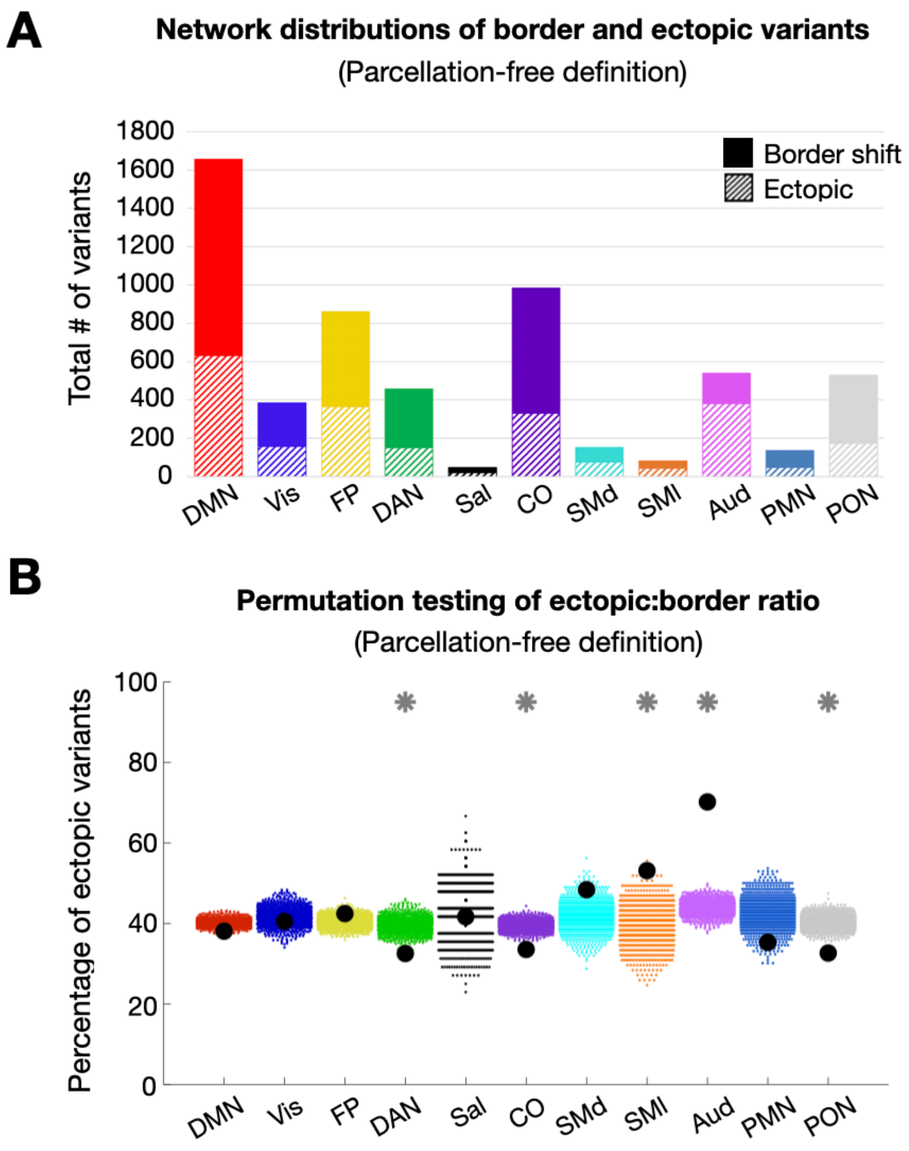


*Supp. Fig. 14: Network linkages of variants using the parcellation-free classification method.* (A) Network assignment distributions of border and ectopic variants in the HCP dataset using the parcellation-free classification method. When implementing the parcellation-free method for classifying border and ectopic variants, network assignment distributions of border vs. ectopic variants are comparable to those obtained using the primary classification method (i.e., compare this figure to Fig. 4A). (B) Permutation testing of ectopic:border ratio of variants classified using the parcellation-free method. While networks such as SMl, Auditory, and PON remain significant compared to the primary method, others exhibit trends in the opposite direction (e.g., CO variants are significantly more likely to be ectopic in the parcellation-free method). Thus, while distributions of border and ectopic classifications are comparable across methods, the significance of their differences is dependent on the analysis method.

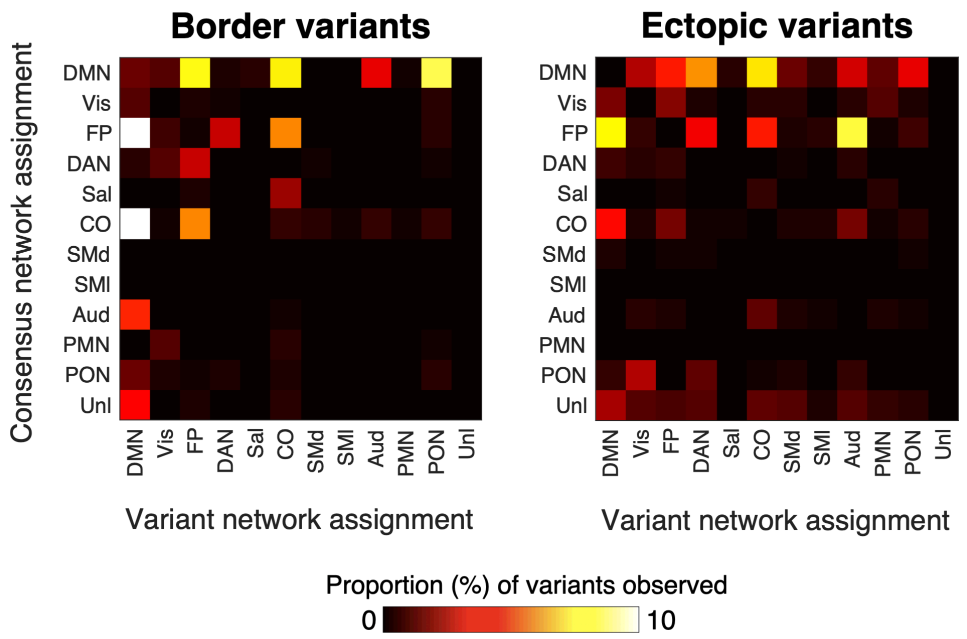


*Supp. Fig. 15: Comparison of variant network assignment to consensus assignments at that location.* For each border (left) and ectopic (right) variant, the figure displays the typical (canonical) network associated with a given variant's location (rows; defined as the modal network across variant vertices) versus the idiosyncratic network to which the variants were assigned in HCP participants (columns). Values represent the raw percentage of variant "swaps" observed (i.e., out of all possible border or ectopic variants). (*Unl.*= network's cortical territory is majority unlabeled in the group-average network description)

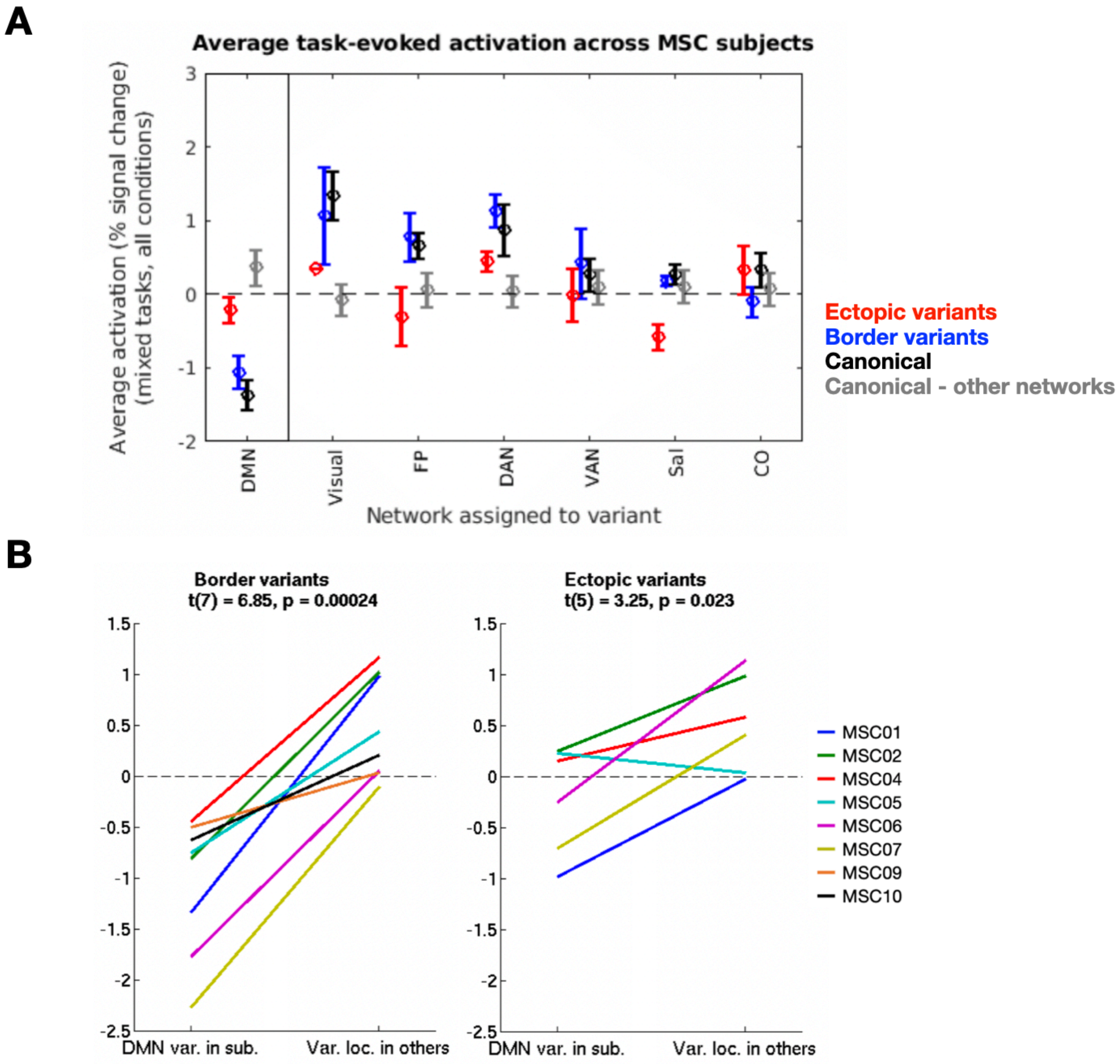


*Supp. Fig. 16: Average task-evoked activation by network across MSC subjects, using the parcellation-free classification method.* (A) Average activation (*z*) across all task conditions in the MSC for variants (red = ectopic, blue = border shift), canonical locations in the listed network (black), or canonical locations in other networks (gray). (B) Average activation of DMN-assigned variants in an individual vs. average activation in the same location in other individuals. Eight of 9 subjects with a border DMN variant and 6 of 9 subjects with an ectopic DMN variant were included. Notably, results are very comparable to those observed using the primary border-ectopic classification method (see Fig. 5 in main manuscript). Colors represent different MSC participants.

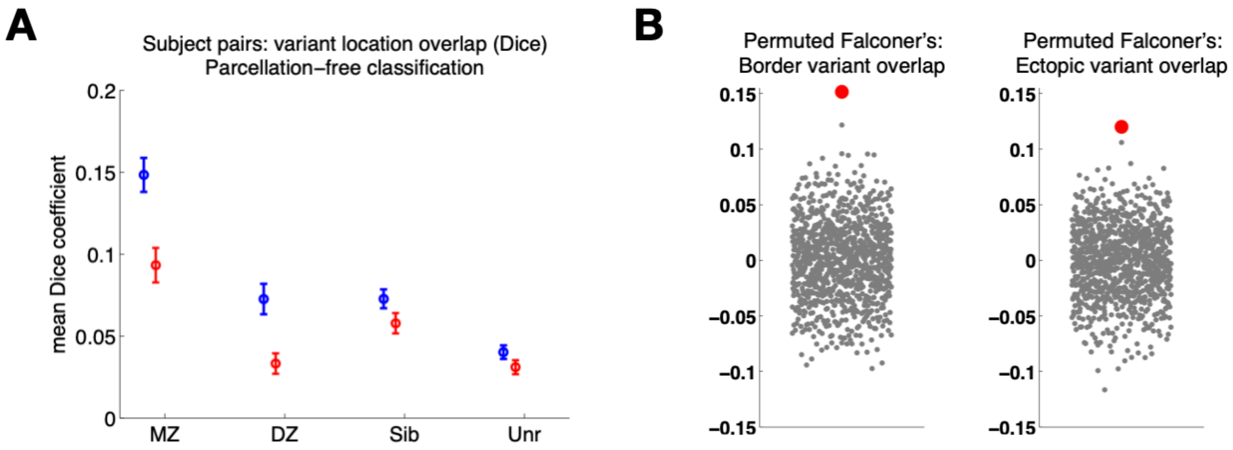


*Supp. Fig. 17: Replication of heritability results using the parcellation-free method to classify variants.* When using a network-independent approach to classify border and ectopic variants, heritability results are replicated; variants are most similar in location among monozygotic twins (exhibiting intermediate similarity among dizygotic twins and siblings), and Falconer's estimates of heritability are significantly higher for both forms of variants relative to a null distribution ($p<0.001$ for both border and ectopic variants).

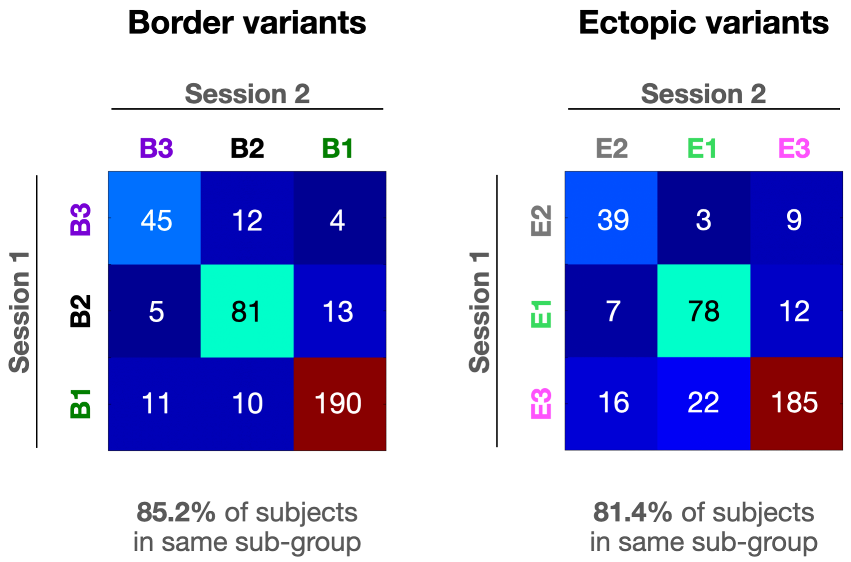


*Supp. Fig. 18: Within-subject consistency of sub-group assignments.* When individuals' data is split by session, most subjects match to the same sub-group for each portion of their data (over 85% of subjects using border variants, and over 81% using ectopic variants; note total $N$ is all 371 subjects with at least one variant of each form). Sub-group profiles used to correlate with individuals' network profiles are those displayed in Fig. 7A/B in the main manuscript (averaged across the two split-halves for each sub-group).

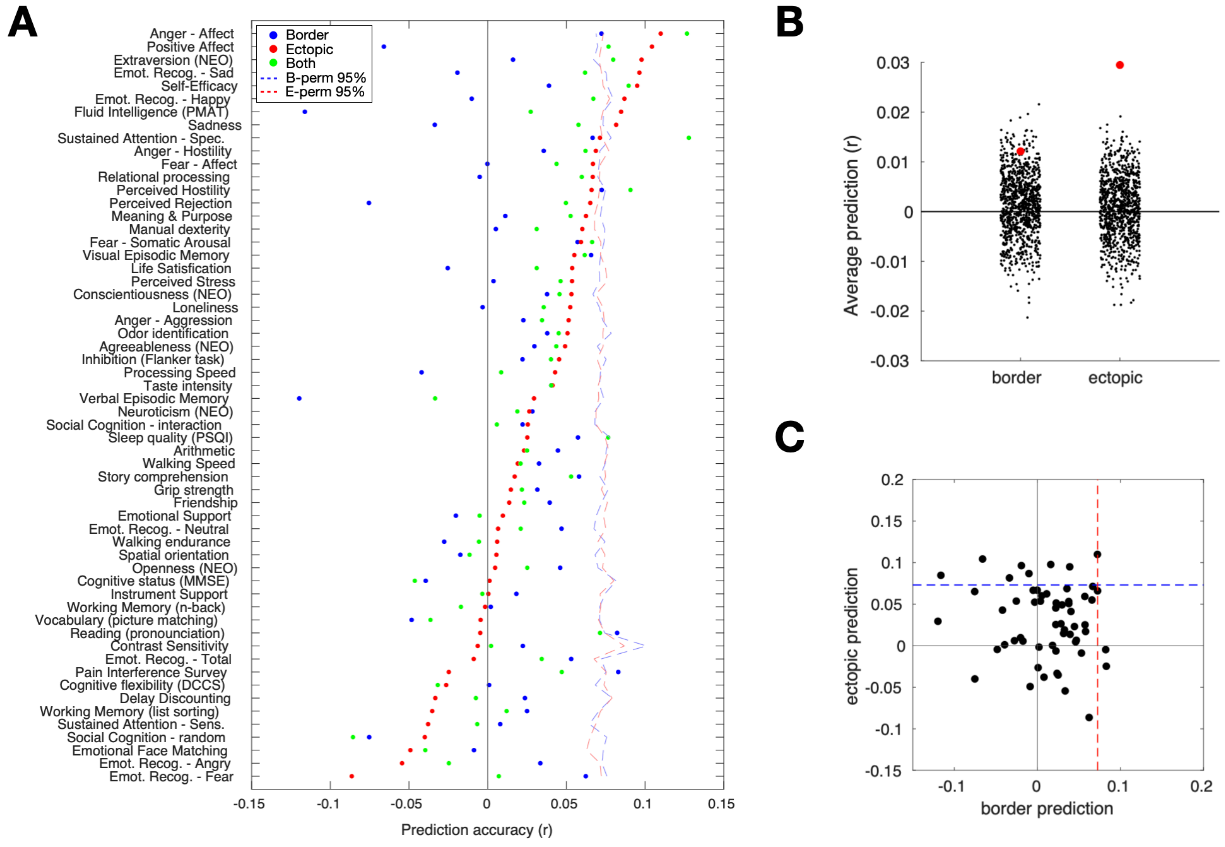


*Supp. Fig. 19: Prediction of behavioral phenotypes based on variant network similarities.* (A) Accuracy for predicting a range of behavioral variables (taken from ref. (Kong et al., 2019)) based on the network affiliations of either ectopic variants (red), border variants (blue), or all variants combined (green); dashed lines = 95% boundary generated from null permutation. (B) Average prediction across all variables for border and ectopic variants (red dots) relative to the average prediction from 1000 null permutations (black dots). Both border (p=0.044) and ectopic (p=0.001) variant features predict behavioral measures to a greater extent than expected by chance. (C) Border and ectopic prediction levels for different variables (black dots) are contrasted with one another directly. Dashed lines represent 95% boundaries from null permutations (blue = border permutations; red = ectopic permutations). Predictions from border and ectopic variants are poorly correlated (r = -0.126), suggesting that they are linked to different behavioral phenotypes.

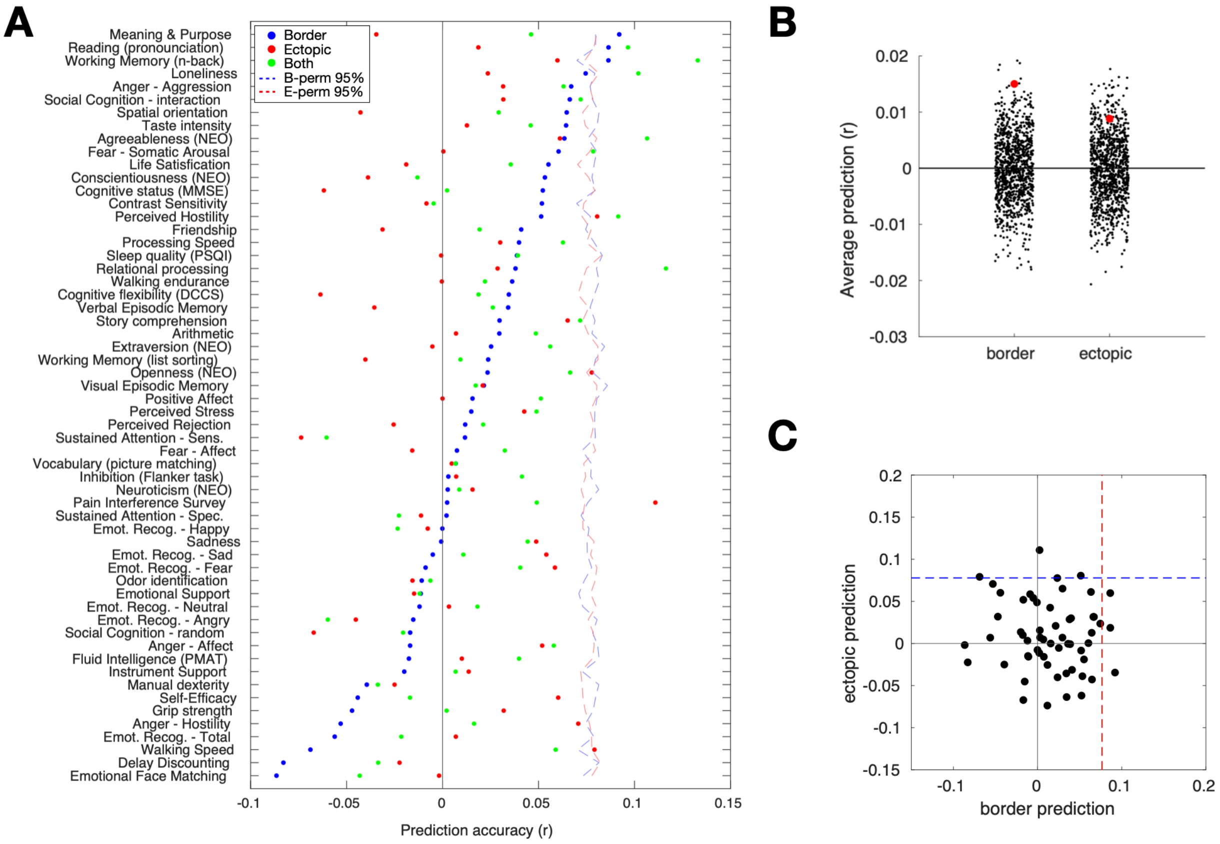


*Supp. Fig. 20: Prediction of behavioral phenotypes based on network variant locations.* (A) Accuracy for predicting a range of behavioral variables (taken from (Kong *et al.*, 2019)) based on the network variant locations of either border (blue) or ectopic (red) variants (dashed lines = 95% boundary generated from null permutation). (B) Average prediction across all variables for border and ectopic variants (red dots) relative to the average prediction from 1000 null permutations (black dots). Only border (p=0.007) variant locations predict behavioral measures on average to a greater extent than expected by chance. (C) Border and ectopic prediction levels for different variables (black dots) are contrasted with one another directly. Dashed lines represent 95% boundaries from null permutations (blue = border permutations; red = ectopic permutations). Predictions from border and ectopic variants are poorly correlated (r = -0.090), suggesting that they are linked to different behavioral phenotypes.

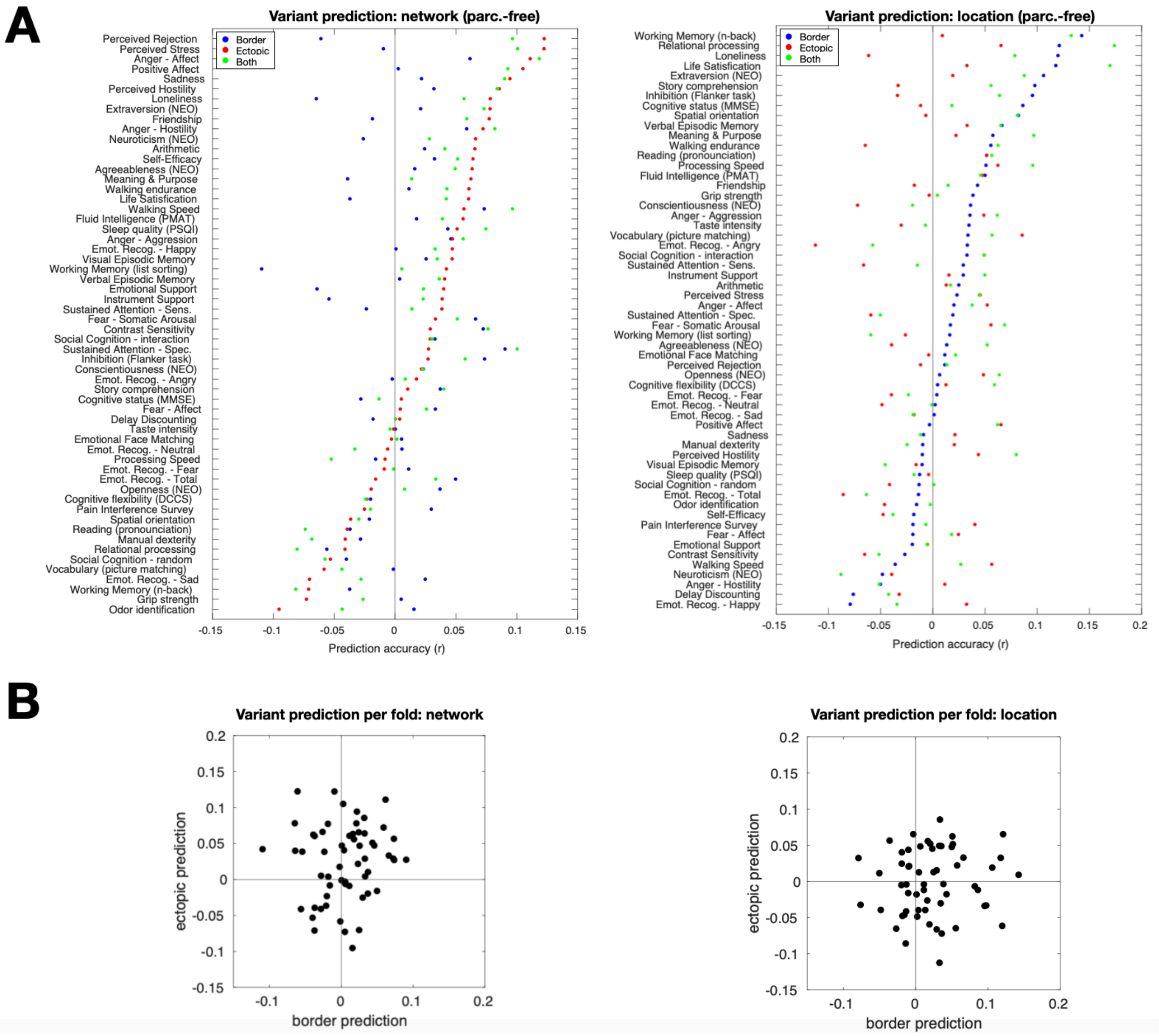


*Supp. Fig. 21: Prediction of behavioral phenotypes based on parcellation-free network similarity measures.* (A) Accuracy for predicting behavioral variables from variant network similarity (left) and variant location (right) of ectopic variants (red), border variants (blue), or all variants combined (green). (B) Border and ectopic prediction levels for different variables (black dots) are contrasted with one another directly based on network (left) and location (right) measures. Predictions from border and ectopic variants are poorly correlated with measures of network similarity (r = 0.09) and variant location (r = 0.073), suggesting that they are linked to different behavioral phenotypes.

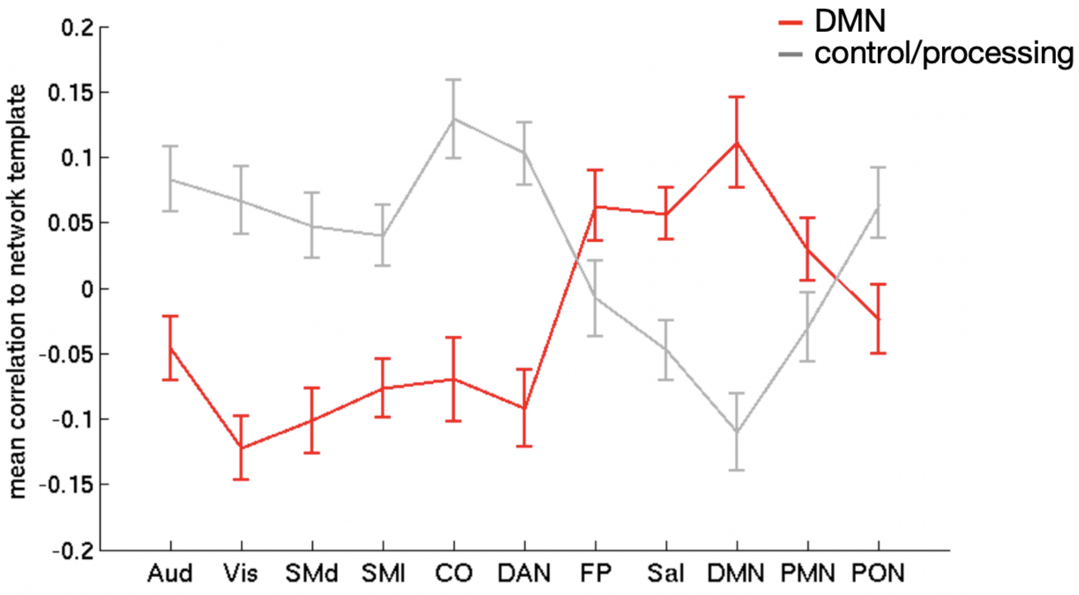


*Supp. Fig. 22: DMN and control/processing sub-group profiles*. In order to determine whether subjects consistently sorted into the same sub-group based on their border vs. ectopic variants, we matched participants into two reliable sub-groups that were identified across all variant forms in previous work (Seitzman *et al.*, 2019): (1) a sub-group of individuals whose variants had high affinity to the DMN and (2) a sub-group of individuals with variants showing higher affinity to top-down control and sensorimotor processing systems. The network profiles for the two subgroups are shown above. These profiles were used as a template to group subjects by both their border and ectopic variants (see *Methods*).

#### SUPPLEMENTAL TABLES

*Supp. Table 1: Ectopic variants from the MSC, classified into different sub-types based on individual brain network comparisons (as described in Supp. Figure 7). Percentages of variants of each sub-type are listed (counts in parentheses), based on three different methods used to classify ectopic variants in the manuscript.*

| Classification method | Islands | Island peaks | Extensions/peninsulas |
|---|---|---|---|
| 3.5mm from network* | 57% (34) | 18% (11) | 25% (15) |
| 10mm from network | 72% (26) | 19% (7) | 8% (3) |
| Parcellation-free[#] | 60% (26) | 19% (8) | 21% (9) |

* Primary method used in paper
### Secondary method (90% of peak at 10mm)

*Supp. Table 2: Contrasts used in analysis of HCP task activations and the tasks to which they belong.*

| **Task** | **Contrast name** |
|---|---|
| Emotion | faces-shapes |
| Emotion | faces |
| Emotion | shapes |
| Gambling | punish-reward |
| Gambling | punish |
| Gambling | reward |
| Language | math-story |
| Language | math |
| Language | story |
| Motor | lf-avg |
| Motor | lh-avg |
| Motor | rf-avg |
| Motor | rh-avg |
| Motor | t-avg |
| Motor | avg |
| Relational | match-rel |
| Relational | match |
| Relational | rel |
| Social | random-tom |
| Social | random |
| Social | tom |
| Working Memory | 2bk-0bk |
| Working Memory | body-avg |
| Working Memory | face-avg |
| Working Memory | place-avg |
| Working Memory | tool-avg |
| Working Memory | 2bk |
| Working Memory | 0bk |

*Supp. Table 3: Variables used for behavioral prediction analysis with official HCP variable names.*

| HCP variable name | Variable description |
|---|---|
| MMSE_Score | Cognitive status (MMSE) |
| PSQI_Score | Sleep quality (PSQI) |
| PicSeq_Unadj | Visual Episodic Memory |
| CardSort_Unadj | Cognitive flexibility (DCCS) |
| Flanker_Unadj | Inhibition (Flanker task) |
| PMAT24_A_CR | Fluid Intelligence (PMAT) |
| ReadEng_Unadj | Reading (pronunciation) |
| PicVocab_Unadj | Vocabulary (picture matching) |
| ProcSpeed_Unadj | Processing Speed |
| DDisc_AUC_40K | Delay Discounting |
| VSPLOT_TC | Spatial orientation |
| SCPT_SEN | Sustained Attention - Sens. |
| SCPT_SPEC | Sustained Attention - Spec. |
| IWRD_TOT | Verbal Episodic Memory |
| ListSort_Unadj | Working Memory (list sorting) |
| ER40_CR | Emot. Recog. - Total |
| ER40ANG | Emot. Recog. - Angry |
| ER40FEAR | Emot. Recog. - Fear |
| ER40HAP | Emot. Recog. - Happy |
| ER40NOE | Emot. Recog. - Neutral |
| ER40SAD | Emot. Recog. - Sad |
| AngAffect_Unadj | Anger - Affect |
| AngHostil_Unadj | Anger - Hostility |
| AngAggr_Unadj | Anger - Aggression |
| FearAffect_Unadj | Fear - Affect |
| FearSomat_Unadj | Fear - Somatic Arousal |
| Sadness_Unadj | Sadness |
| LifeSatisf_Unadj | Life Satisfaction |
| MeanPurp_Unadj | Meaning & Purpose |
| PosAffect_Unadj | Positive Affect |
| Friendship_Unadj | Friendship |
| Loneliness_Unadj | Loneliness |
| PercHostil_Unadj | Perceived Hostility |
| PercReject_Unadj | Perceived Rejection |
| EmotSupp_Unadj | Emotional Support |

| | |
|---|---|
| InstruSupp_Unadj | Instrument Support |
| PercStress_Unadj | Perceived Stress |
| SelfEff_Unadj | Self-Efficacy |
| Emotion_Task_Face_Acc | Emotional Face Matching |
| Language_Task_Story_Avg_Difficulty_Level | Story comprehension |
| Language_Task_Math_Avg_Difficulty_Level | Arithmetic |
| Relational_Task_Acc | Relational processing |
| Social_Task_Perc_Random | Social Cognition - random |
| Social_Task_Perc_TOM | Social Cognition - interaction |
| WM_Task_Acc | Working Memory (n-back) |
| Endurance_Unadj | Walking endurance |
| GaitSpeed_Comp | Walking Speed |
| Dexterity_Unadj | Manual dexterity |
| Strength_Unadj | Grip strength |
| NEOFAC_A | Agreeableness (NEO) |
| NEOFAC_O | Openness (NEO) |
| NEOFAC_C | Conscientiousness (NEO) |
| NEOFAC_N | Neuroticism (NEO) |
| NEOFAC_E | Extraversion (NEO) |
| Odor_Unadj | Odor identification |
| PainInterf_Tscore | Pain Interference Survey |
| Taste_Unadj | Taste intensity |
| Mars_Final | Contrast Sensitivity |
